# TAR syndrome causal gene *RBM8A* is critical for embryonic bone development and proper Hedgehog signaling

**DOI:** 10.64898/2026.04.21.718480

**Authors:** Jennifer Mott, Zifei Pei, Jiawan Sun, Choong Heon Lee, Twishi Puri, Zander M. Sachs, Ana Aviles Vargas, Nicholas Blaha, Mabel Tong, Aimin Liu, Yong Wang, Yongsoo Kim, Santhosh Girirajan, Jiangyang Zhang, Yingwei Mao

## Abstract

RNA-Binding Motif Protein 8A (RBM8A) is a core component of the exon junction complex (EJC), which plays a fundamental role in post-transcriptional RNA regulation. Mutations in the *RBM8A* gene cause thrombocytopenia-absent radius (TAR) syndrome, characterized by reduced platelet number and radial defects. However, how its function contributes to bone development remains poorly understood. We found that RBM8A is highly expressed in human bone marrow-derived mesenchymal stem cells (MSCs) and mouse embryonic limb buds. To further investigate the role of RBM8A in bone development, we generated an *Rbm8a* conditional knockout mouse (cKO) model in the early limb bud and craniofacial mesenchyme using the *Prx1-cre* line, which recapitulates key features of TAR syndrome. *Rbm8a* deficiency in this mouse model resulted in significant defects in limb and craniofacial development, including shorter limb bones across developmental timepoints, delayed ossification, altered cranial morphogenesis, along with impairments in motor function and reduced brain size. To elucidate the molecular basis of these phenotypes, we performed RNA immunoprecipitation sequencing (RIP-seq) in embryonic limb tissue. Our analysis revealed that RBM8A preferentially associates with long, exon-rich transcripts with high isoform complexity. These transcripts are enriched in pathways related to primary cilia, RNA processing, and developmental signaling. Consistent with this, transcriptomic and enrichment analyses identified strong associations with ciliopathy-related and congenital skeletal disease gene sets. At the cellular level, *Rbm8a* deficiency resulted in altered expression of key chondrogenic and ossification markers in embryonic limb bone tissue, such as reduced SOX9 and increased IHH in the primary ossification center, suggesting impaired progression of endochondral ossification and disrupted Hedgehog pathway activity. Quantitative Reverse Transcription–PCR (qRT-PCR) analysis further validated selective downregulation of Hedgehog pathway targets, including *Gli3* and *Hhip*, while other developmental genes remained unchanged, indicating pathway-specific attenuation. Together, these findings support a model in which *Rbm8a* regulates structurally complex transcripts that are enriched in ciliary and developmental signaling pathways, and that disruption of this process leads to selective impairment of Hedgehog signaling and skeletal development. These results provide a novel mechanistic link between RNA processing and developmental signaling and offer new insight into how defects in RNA regulation contribute to congenital skeletal abnormalities.

## Introduction

Birth defects impacting skeletal and craniofacial development are among the most common congenital diseases, representing some of the urgent unmet medical needs across the world^1^. The prevalence of major structural birth defects, including skeletal and craniofacial abnormalities, is alarmingly high, affecting 3% of newborns in the US, with an annual hospitalization cost of $22.6 billion in 2013^2^. Limb reduction defects (LRD) and craniofacial anomalies (CFA) are particularly common, affecting 1 in 1900 and 1 in 1600 babies, respectively^3^. Of note, congenital bone disorders represent a major group of developmental conditions that cause significant lifelong structural and functional health complications^4^. Moreover, these conditions are often associated with nervous system defects, such as autism spectrum disorders (ASD) and intellectual disability (ID)^5–8^. Therefore, there is a significant medical need for individuals affected by these debilitating disorders.

These diseases are caused by genetic or environmental factors, and identifying the direct causal factor(s) for each disorder is critical to understanding fundamental mechanisms of bone development. Skeletal development is a highly coordinated process that begins during early embryogenesis, progressing through tissue patterning, mesenchymal condensation, chondrogenesis, endochondral ossification, growth plate maturation, and continuing into postnatal remodeling^9^. Because skeletal fate is established early during embryogenesis, it is uniquely sensitive to disruptions in developmental signaling^10^. Bone morphogenesis is precisely arranged by interconnected signaling pathways that control cellular proliferation, lineage determination, and maturation timing^11^. Hedgehog signaling is one such major pathway, as Sonic Hedgehog (SHH) governs early limb patterning and Indian Hedgehog (IHH) controls endochondral ossification^12^. As such, IHH mutations cause congenital skeletal disorders such as Brachydactyly Type A1 or Acrocapitofemoral dysplasia^13,14^. Moreover, mutations in GLI3, a central regulator of Hedgehog signaling, result in congenital limb and craniofacial disorders such as Pallister–Hall syndrome or Greig cephalopolysyndactyly syndrome^15,16^. Investigating the developmental mechanisms that give rise to specific bone phenotypes can provide insights into core signaling pathways that are critical during early development.

Although more genetic associations are being identified, the mechanisms by which these genes affect bone development remain poorly understood^4,11,17^. One example is TAR syndrome, which is a rare congenital disorder characterized by the absence of radius in each forearm (with the presence of both thumbs) along with a deficiency in platelet counts (less than 50 platelets/nL, where the normal range is 150-400 platelets/nL)^18^. In addition to these anomalies, the severity of other skeletal abnormalities varies among those with this syndrome^19^.

Craniofacial and other upper- and lower-limb bones can also be underdeveloped, affecting mobility and other motor functions involved in everyday tasks for TAR patients^20^. Moreover, some TAR patients have hypoplasia of the cerebellar vermis and the corpus callosum^21^. This brain underdevelopment, along with the prevalence of cranial defects and intellectual disability^19^, suggests there may be distinct changes in the crania and brains of affected TAR patients.

The causal factors of TAR syndrome are compound mutations of a 1q21.1 null deletion and rare single-nucleotide polymorphisms in the regulatory region of *RBM8A*^22^. *RBM8A*, also known as Y14, is a core EJC factor, which is critical for splicing, translation, and mRNA degradation^23^. EJC plays a major role in nonsense-mediated mRNA decay (NMD), which controls RNA stability by tagging mRNAs with premature termination codons for decay^23^. *RBM8A* has previously been reported to regulate cell cycle progression and is implicated in cancer, neurodegenerative diseases, and neurodevelopmental disorders^24^. Previous works from our group^25–28^ and others^29^ have demonstrated the critical role of *RBM8A* in brain development. However, the role of *RBM8A* in bone development and the mechanisms by which its mutations cause the specific phenotypic anomalies in TAR syndrome remain unclear^30^. Understanding how *RBM8A* gene mediates embryonic bone development is critical not only for advancing knowledge of congenital skeletal disorders, but also for identifying potential therapeutic targets and uncovering new roles these genes may play in key developmental signaling pathways.

To address this knowledge gap, we present a novel *Rbm8a* cKO mouse model, which resembles the TAR phenotype. We demonstrate that these mice exhibit structural skeletal defects across developmental timepoints, functional impairments, and changes in the brain. Importantly, we identify the altered Hedgehog signaling pathway as the mechanism by which loss of *RBM8A* leads to limb maldevelopment in TAR syndrome. These findings will serve as a basis for understanding the novel role of RBM8A in skeletal development. Insights into *RBM8A*’s functions in long bone and cranial development, and its implication in brain development, will be the first critical step in therapeutic development for this disorder.

## Results

### Expression of RBM8A in developing bones

As *RBM8A* deficiency leads to TAR syndrome with well-defined bone defects, we first examined the RBM8A level in bone-related cells and tissues. MSCs, characterized by surface markers such as CD90, can differentiate into osteogenic cell types^31^. We cultured human bone marrow-derived MSCs and found that RBM8A is highly expressed in CD90-positive cells (Figure 1A).

**Figure 1:**
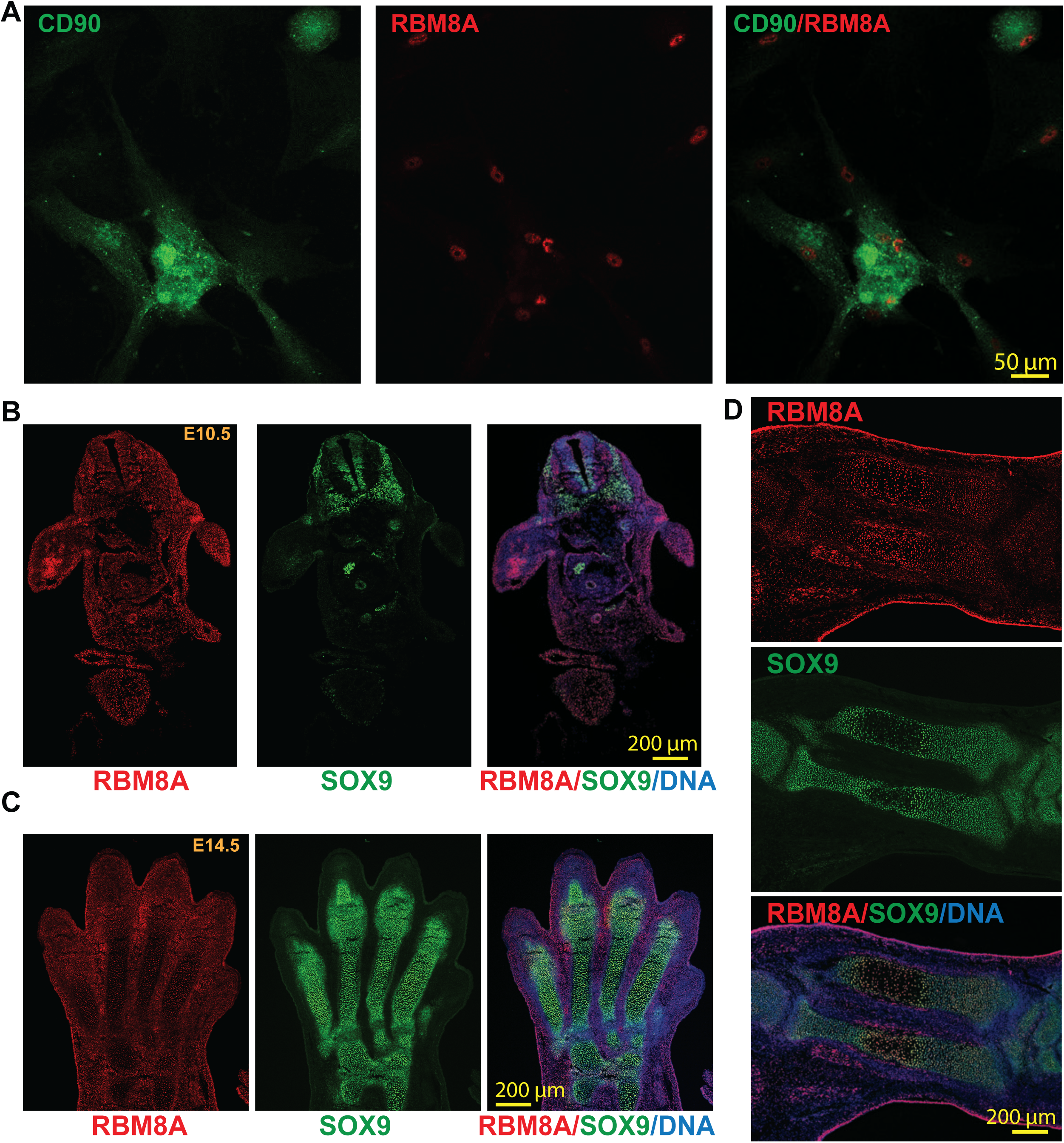
RBM8A is expressed in human bone marrow-derived mesenchymal stem cells and mouse embryonic skeletal tissues. **A.** Immunofluorescence staining of cultured human bone marrow-derived mesenchymal stem cells showing CD90 (green) and RBM8A (red). Merged image (right) demonstrates RBM8A localization in CD90-positive cells. Scale bar = 50 µm. **B.** Whole-mount immunofluorescence staining of E10.5 mouse embryo section showing RBM8A (red), SOX9 (green), and merged image with DAPI (blue). RBM8A is broadly expressed, with high expression in the limb bud. In the neural crest, RBM8A expression is decreased in SOX9-positive regions. Scale bar = 200 µm. **C.** Whole-mount immunofluorescence staining of E14.5 mouse forelimb autopod sections showing RBM8A (red), SOX9 (green), and merged image with DAPI (blue). RBM8A is broadly distributed throughout the autopod, including SOX9-positive cartilage condensation. Scale bar = 200 µm. **D.** Whole-mount immunofluorescence staining of E14.5 mouse forelimb zeugopod longitudinal tissue section showing RBM8A (red), SOX9 (green), and merged image with DAPI (blue). RBM8A is detected in and around SOX9-positive skeletal condensations, including the central region of the developing cartilage element. Scale bar = 200 µm.

Moreover, to determine whether RBM8A is expressed in the early embryonic limb bud, we performed co-immunostaining for RBM8A and SOX9, a key regulator of pre-chondrogenic specification^32^, in embryonic day 10.5 (E10.5) mouse sections. We found that RBM8A is ubiquitously expressed in most cells, with higher expression in the limb bud (Figure 1B).

Interestingly, RBM8A expression is decreased in SOX9-positive cells compared with SOX9-negative neural crest cells. We further examined RBM8A and SOX9 expression in both embryonic autopods and zeugopods (radius and ulna) at E14.5 (Figure 1C-D). SOX9-positive chondroprogenitors and chondrocytes are localized in the digital cartilage condensations within the autopod but are absent in the interdigital mesenchymal cells (Figure 1C). In contrast, RBM8A expression is higher in the interdigital mesenchymal cells that are SOX9-negative but lower in SOX9-positive cells. Consistently, in both the radius and ulna, RBM8A expression is higher in SOX9-negative hypertrophic chondrocytes in the center of the diaphysis than in SOX9-positive chondrocytes at the sides of the diaphysis (Figure 1D), suggesting an inverse relationship between SOX9 and RBM8A. These data collectively suggest that RBM8A could play an essential role in early forelimb development.

### Generation of *Prx1-Rbm8a^f/+^* mice

To study the role of RBM8A in embryonic limb development, we generated an *Rbm8a* limb cKO mouse model. We used *Prx1-Cre* mice to drive the expression of Cre recombinase throughout the early limb bud mesenchyme and a subset of craniofacial mesenchyme^33^ (Figure 2A; Supplementary Fig. 1A). The resulting *Prx1-Rbm8a^f/+^* mice have noticeably shorter forelimbs and forepaws, as well as significantly smaller overall body length compared to their littermate controls (Figure 2B). Quantification of Alizarin Red and Alcian Blue staining revealed significantly shorter digit, radius, and humerus lengths in *Prx1-Rbm8a^f/+^* mice compared to their littermate controls (Figure 2C, 2D). This phenotype is observable as early as E15 (Figure 2E), in which the *Prx1-Rbm8a^f/+^* embryos show notably shorter limbs and altered cranial shape.

**Figure 2:**
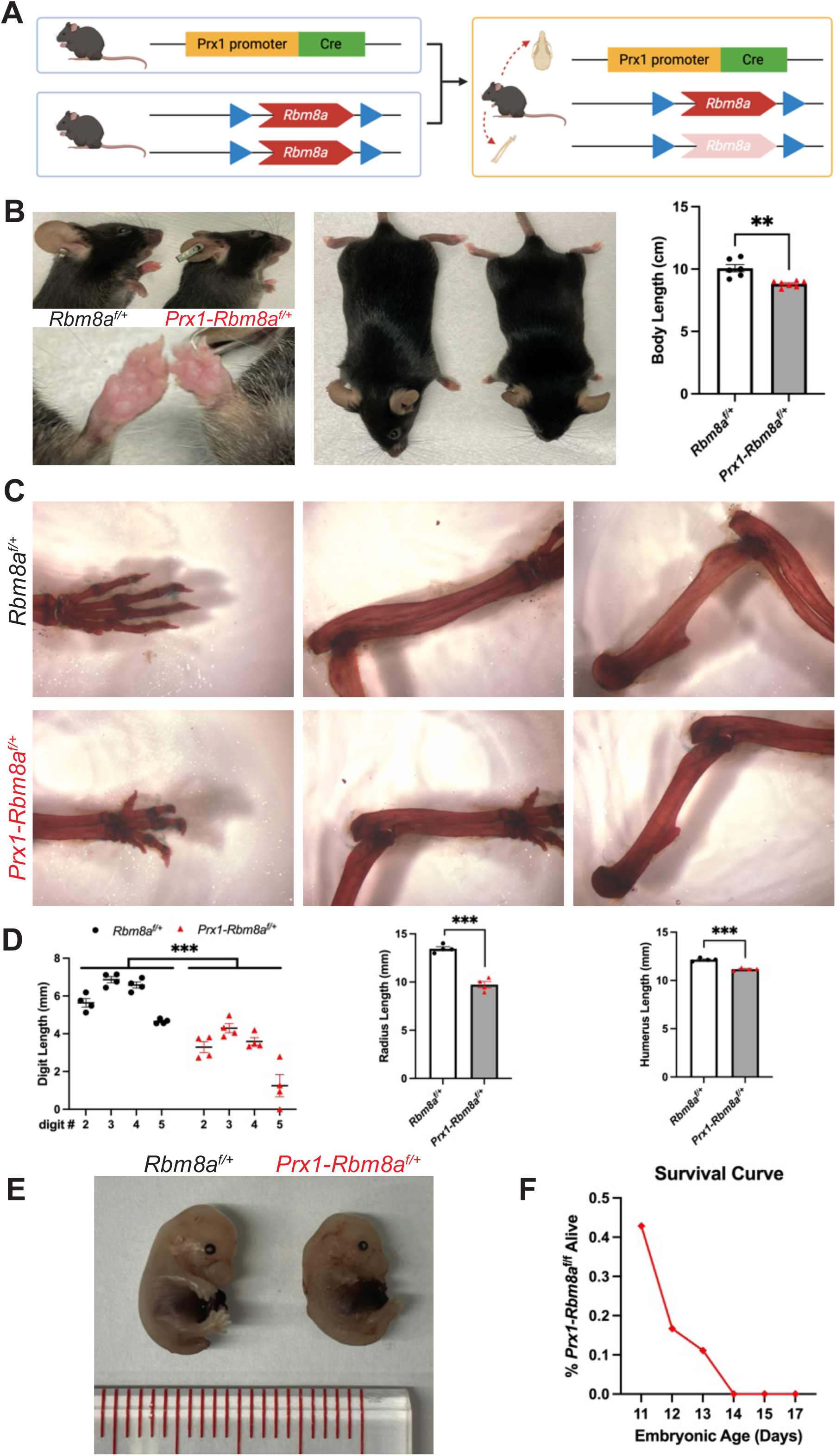
*Prx1-Rbm8a^f/+^* mice show limb and craniofacial malformation. **A.** Schematic of the breeding strategy used to generate *Prx1-Rbm8a^f/+^* mice. *Rbm8a^f/f^* mice were crossed with *Prx1-Cre* mice to generate *Prx1-Cre; Rbm8a^f/+^* offspring, which have selective deletion of *Rbm8a* in limb bud mesenchyme and some craniofacial mesenchyme. Created with BioRender.com. **B.** Overall phenotype of *Prx1-Rbm8a^f/+^* mice (right) compared to sex-matched littermate control (left). *Prx1-Rbm8a^f/+^* mice have substantially shorter forelimbs and forepaws, and significantly smaller overall body length. Mean ± SEM (n=6 per group). Unpaired *t*-test. **, p < 0.01. **C.** Representative images of Alizarin Red and Alcian Blue staining of forelimb bones (left: forepaws, center: radius and ulna, right: humerus) of adult *Prx1-Rbm8a^f/+^* mice (bottom) and sex-matched littermate control (top). **D.** Quantification and statistical analysis of Alizarin Red and Alcian Blue staining. **Left:** all digits measured (II to V) of *Prx1-Rbm8a^f/+^* mice were significantly shorter than those of their littermate controls. **Middle:** radii of *Prx1-Rbm8a^f/+^* mice were significantly shorter than those of their littermate controls. **Right:** humeri of *Prx1-Rbm8a^f/+^* mice were significantly shorter than those of their littermate controls. Mean ± SEM (n=4 per group). Unpaired *t*-test. ***, p < 0.001. **E.** Comparison of *Prx1-Rbm8a^f/+^* (right) and control (left) E15 embryos collected from the same litter. The *Prx1-Rbm8a^f/+^*embryo shows noticeably smaller body, shorter limbs, and an altered craniofacial region compared to the control. **F.** Survival curve of homozygous deletion *Prx1-Rbm8a^f/f^* embryos. At E11, 42.9% of *Prx1-Rbm8a^f/f^* embryos were viable, decreasing to 16.7% at E12 and 11.1% at E13, with no viable embryos detected from E14 onward.

Forelimb and cranial Ai14 reporter signal intensities of optically cleared *Prx1-Rbm8a ^f/+^; Ai14 ^f/+^* E15 embryos were significantly reduced than those of *Prx1; Ai14 ^f/+^* (control) (Supplementary Fig. 1). Interestingly, we observed embryonic lethality in homozygous deletion *Prx1-Rbm8a^f/f^*mice. At E11, 42.9% of *Prx1-Rbm8a^f/f^* embryos were alive. The survival rate decreased sharply to 16.7% at E12, then to 11.1% at E13, eventually reaching 0% from E14 onward (Figure 2F). This indicates that *Rbm8a* is required for the *Prx1-cre* mesenchymal lineage during early development. In all, our *Prx1-Rbm8a^f/+^* mouse model captures the limb and craniofacial malformation observed in patients with TAR syndrome.

*Prx1-Rbm8a^f/+^* mice show significant bone defects across developmental timepoints To further investigate the alterations in bone development, we utilized micro-computed tomography (micro-CT) to measure the lengths of the limb bones in *Prx1-Rbm8a^f/+^* mice and control littermates at postnatal days 5, 15, and 30 (P5, P15, and P30). The stark difference in bone phenotype can be observed via micro-CT as early as at P5 and becomes increasingly noticeable as they develop (Figure 3). At P5, control mice show expected early postnatal mineralization^34,35^, whereas *Prx1-Rbm8a^f/+^* mice already show shorter long bones and delayed cranial growth (Figure 3A). At P15, which is an active skeletal growth phase, control mice demonstrate an overall increase in bone length and stronger cranial mineralization than at P5^34^ (Figure 3B). Contrastingly, *Prx1-Rbm8a^f/+^*mice continue to display shortened long bones and delayed cranial maturation (Figure 3B). At P30, control mice show expected near-mature skeletal growth^34,36^, but *Prx1-Rbm8a^f/+^* mice have persistently reduced overall bone length, indicating long-term structural developmental consequences (Figure 3C). When quantified, we saw statistically significantly shorter lengths of radius, ulna, humerus, tibia, and femur bones in the *Prx1-Rbm8a^f/+^* mice compared to their littermate controls across all three ages (Figure 3D-3H). Interestingly, the radius and ulna lengths of *Prx1-Rbm8a^f/+^* mice appear to halt between P15 and P30 (Figure 3D, 3E), suggesting that there is a defect in postnatal longitudinal bone growth between these developmental timepoints.

**Figure 3:**
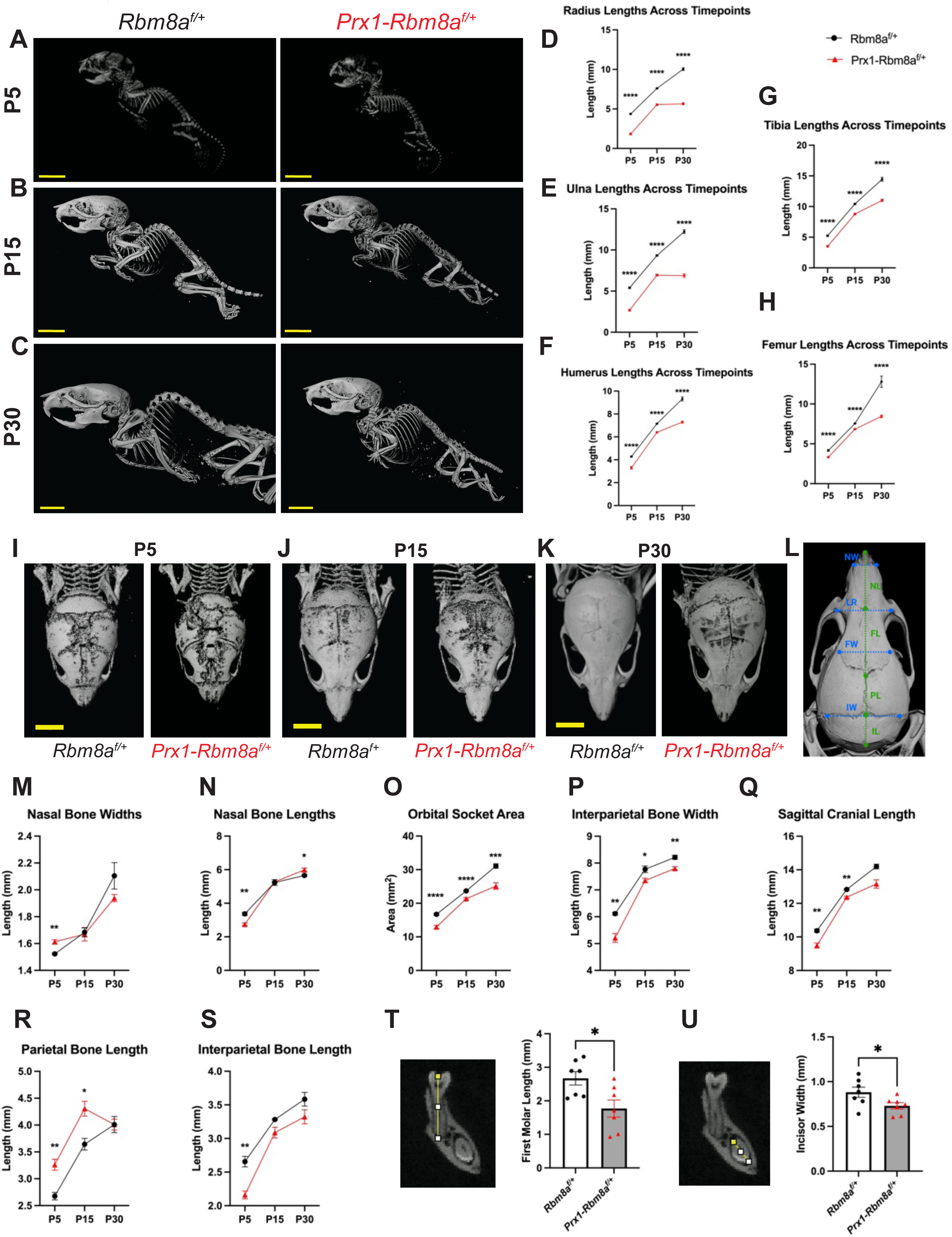
Micro-CT of *Prx1-Rbm8a^f/+^* mice shows significant bone defects across developmental timepoints. **A-C.** Representative whole body images of micro-CT scans of (A) P5, (B) P15, (C) P30 *Prx1-Rbm8a^f/+^* mice (right) and control mice (left). Scale bar = 7mm. **D-H.** Quantification and statistical analyses of (D) radius, (E) ulna, (F) humerus, (G) tibia, (H) femur lengths of *Prx1-Rbm8a^f/+^*mice (red) and control mice (black) across developmental timepoints (P5, P15, P30). Mean ± SEM (n=8 per group). Unpaired *t*-test. ****, p < 0.0001. **I-K.** Representative micro-CT scan image of crania of (I) P5, (J) P15, (K) P30 *Prx1-Rbm8a^f/+^* mice (right) and control mice (left). Scale bar: 6mm. **L.** Cranial landmarks used for quantification. NW: nasal bone width, NL: nasal bone length, LR: anterolateral distance, FW: frontal bone width, FL: frontal bone length, PL: parietal bone length, IW: interparietal bone width, IL: interparietal bone length. **M-S.** Quantification and statistical analyses of (M) nasal bone widths, (N) nasal bone lengths, (O) orbital socket area, (P) parietal bone length, (Q) interparietal bone width, (R) interparietal bone width, (S) sagittal cranial length of *Prx1-Rbm8a^f/+^* mice (red) and control mice (black) across developmental timepoints (P5, P15, P30). Mean ± SEM (n=3 per group). Unpaired *t*-test. **, p < 0.01. **T.** Representative micro-CT scan image of first molars (left) and quantification and statistical analyses (right) of first molar length of *Prx1-Rbm8a^f/+^* mice and control mice. Mean ± SEM (n=7). Unpaired *t*-test. *, p < 0.05. **U.** Representative micro-CT scan image of incisors (left) and quantification and statistical analyses (right) of incisor widths of *Prx1-Rbm8a^f/+^* mice and control mice. Mean ± SEM (n=7). Unpaired *t*-test. *, p < 0.05.

In addition to the limb bones, we also scanned the crania of *Prx1-Rbm8a^f/+^*mice and control littermates at P5, P15, and P30 (Figure 3I-3K). At P5, all mice show partially mineralized frontal and parietal bones and open sutures^37^, but *Prx1-Rbm8a^f/+^* mice show more drastic underdevelopment of nasal, frontal, parietal, and interparietal bones compared to those of littermate mice (Figure 3I). At P15, the control mice show expected increased mineralization of frontal and parietal bones and narrowing sutures than at P5^37^ (Figure 3J). Conversely, the crania of *Prx1-Rbm8a^f/+^*mice continue to show less mineralization and underdevelopment of sutures, especially in the frontal and parietal region, compared to those of littermate mice (Figure 3J). At P30, the control mice show structurally adult-like crania^37^, but *Prx1-Rbm8a^f/+^* mice persistently display reduced mineralization, as well as crooked sutures compared to those of littermate mice (Figure 3K). This misalignment may indicate the loss of coordination and balance between osteoblast and suture mesenchyme activity^38,39^.

As patients with TAR syndrome are often present with abnormal head size^40^, we next measured specific cranial bones of *Prx1-Rbm8a^f/+^*mice and control littermates at P5, P15, and P30 using established cranial landmarks^37,41,42^ (Figure 3L). Interestingly, *Prx1-Rbm8a^f/+^*mice had significantly wider nasal bones than littermates at P5, but showed no statistical difference at P15, and then showed a trend toward being narrower at P30 (Figure 3M). Similarly, they had significantly shorter nasal bones at P5, showed no statistical difference at P15, and exhibited a trend toward being slightly longer at P30 (Figure 3N). These results seem to suggest a dynamic bone growth event that is defective in *Prx1-Rbm8a^f/+^* mice. Orbital socket area, interparietal bone width, and sagittal cranial length were significantly smaller in *Prx1-Rbm8a^f/+^* mice across all three ages (Figure 3O-3Q; Supplementary Fig. 2E, 2F). Notably, *Prx1-Rbm8a^f/+^* mice had significantly longer parietal bones than littermates at P5 and P15 but had no statistical difference at P30 (Figure 3R). These mice also had significantly shorter interparietal bones than their littermates at P5, but there was no statistical difference at P15 or P30 (Figure 3S). Additionally, their frontal bones were significantly narrower than those of their littermates at P30 (Supplementary Fig. 2A). Intriguingly, these bones were significantly shorter than those of their littermates at P5, but caught up by P15, only to be significantly shorter comparatively again at P30 (Supplementary Fig. 2B). The distance between the anterolateral corners of the frontal bone and the height of cranial cavity between bregma and intersphenoidal synchondrosis were comparable between *Prx1-Rbm8a^f/+^* and control mice across all three timepoints (Supplementary Fig. 2C, 2D). To further examine the craniofacial differences, we also measured the teeth of these mice^43^. Interestingly, *Prx1-Rbm8a^f/+^* mice had significantly shorter first molars and significantly narrower incisors (with no difference in length) compared to littermate controls (Figure 3T, 3U; Supplementary Fig. 2G). Taken together, these measurements reflect an altered craniofacial development in *Prx1-Rbm8a^f/+^* mice.

### *Prx1-Rbm8a* cKO mice show significant motor function deficits and potentially altered psychiatric states

Building on our observations of the striking structural phenotype, we wanted to test whether the structural limb defects also affect functional abilities in *Prx1-Rbm8a^f/+^*mice. First, we used a grip-strength meter to measure the maximum force exerted by the forelimbs of the mice^44,45^.

*Prx1-Rbm8a^f/+^* mice showed significantly weaker grip strength (normalized by body weight) compared to control mice (Figure 4A). Next, we used a slightly elevated apparatus with a wire grid to test the number of paw slips through the wire^46,47^. *Prx1-Rbm8a^f/+^* mice had a significantly higher number of foot slips compared to their counterparts, indicating a motor coordination deficit (Figure 4B). Similarly, *Prx1-Rbm8a^f/+^*mice fell from the rotarod significantly faster than control mice across all three days of testing (Figure 4C). Notably, both control and *Prx1-Rbm8a^f/+^* mice took significantly longer to fall across the days, indicating that motor skill learning and memory were not affected^48,49^ (Figure 4C). Additionally, *Prx1-Rbm8a^f/+^*mice buried significantly fewer marbles than control mice in the marble burying test (Figure 4D), further suggesting fine motor impairment^50^. In the open field test^51^, control and *Prx1-Rbm8a^f/+^*mice traveled similar total distances, but *Prx1-Rbm8a^f/+^* mice spent significantly less time not moving than control mice (Figure 4E). This is interesting because it suggests that *Prx1-Rbm8a^f/+^* mice exhibited higher locomotor activity despite their shortened limb phenotype and apparent motor impairment. This can be interpreted as *Prx1-Rbm8a^f/+^*mice exhibiting hyperactivity or heightened arousal^52^. In addition, *Prx1-Rbm8a^f/+^* mice spent significantly more time in the border than control mice (Figure 4E), which can also suggest increased anxiety-like behavior^52^. There was no significant difference observed in the average velocity, latency to the center, or frequency in the center (Supplementary Fig. 3A). In the elevated plus maze^53^, *Prx1-Rbm8a^f/+^*mice spent significantly less time not moving than control mice, although there was no statistically significant difference in the total distance traveled, time spent in or latency to open arms, frequency in open arms, or average velocity (Figure 4F; Supplementary Fig. 3B).

**Figure 4:**
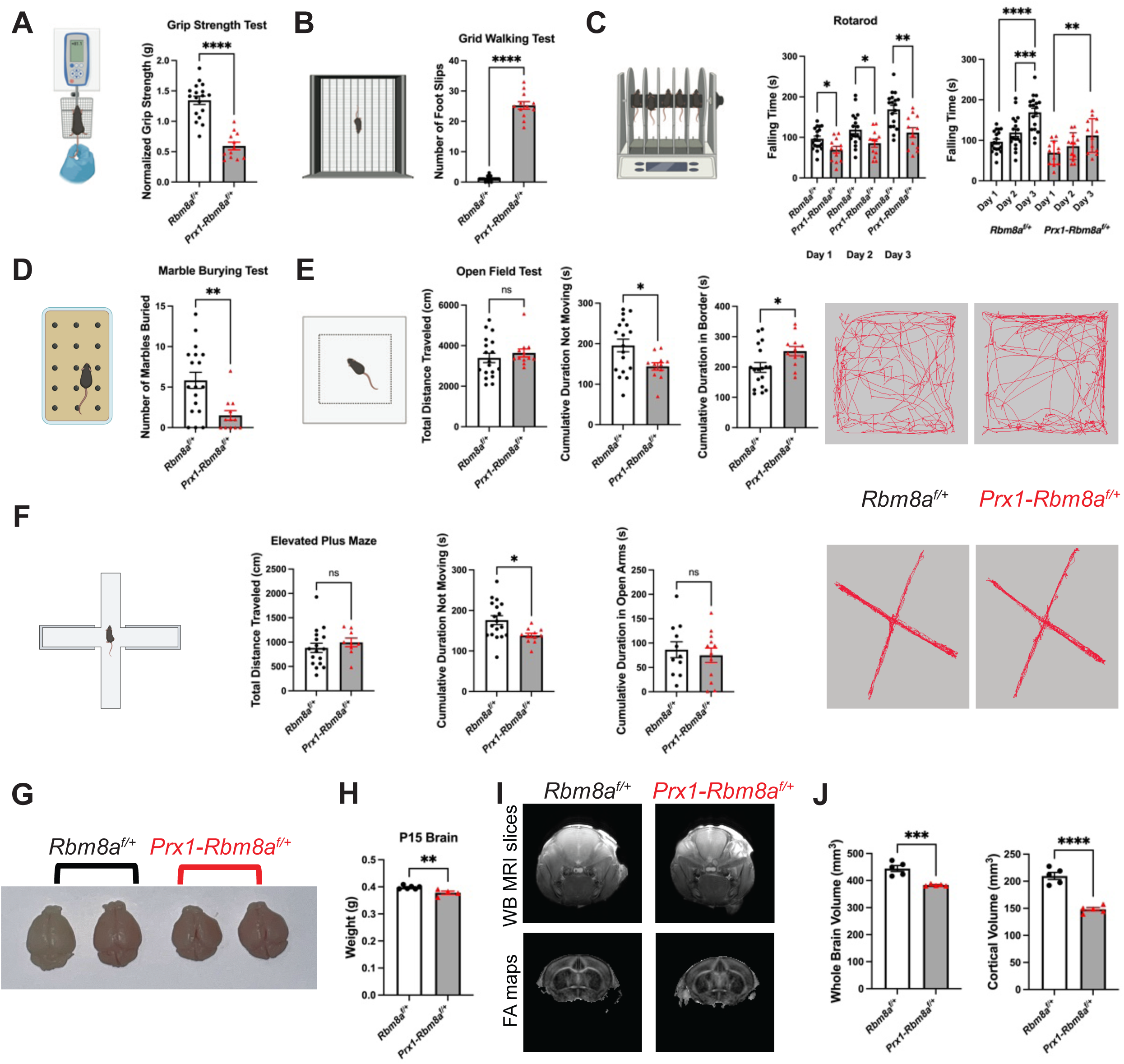
*Prx1-Rbm8a^f/+^* mice show significant motor function deficits and potentially altered psychiatric states. **A.** Grip strength test to measure the maximum force exerted by the forelimbs of the mice; normalized by body weight. *Prx1-Rbm8a^f/+^* mice show significantly weaker grip strength compared to control mice. Mean ± SEM (n=30 total; control: n=18 (9 males, 9 females); mutant: n=12 (5 males, 7 females)). Unpaired *t*-test. ****, p < 0.0001. Created with BioRender.com. **B.** Grid walking test to measure motor coordination function. *Prx1-Rbm8a^f/+^* mice show a significantly higher number of paw slips through the wire grid compared to control mice. Mean ± SEM (n=30 total; control: n=18 (9 males, 9 females); mutant: n=12 (5 males, 7 females)). Unpaired *t*-test. ****, p < 0.0001. Created with BioRender.com. **C.** Rotarod test to measure motor coordination and motor learning across three days. *Prx1-Rbm8a^f/+^* mice show significantly shorter time to fall from the rotarod than control mice across all three days of testing. Both control and *Prx1-Rbm8a^f/+^* mice took significantly longer to fall from the rotarod by day 3 compared to day 1. Mean ± SEM (n=30 total; control: n=18 (9 males, 9 females); mutant: n=12 (5 males, 7 females)). Unpaired *t*-test. *, p < 0.05, **, p < 0.01, ***, p < 0.001, ****, p < 0.0001. Created with BioRender.com. **D.** Marble burying test to measure fine motor coordination of the forepaw and forelimb. *Prx1-Rbm8a^f/+^* mice buried significantly fewer marbles than control mice, suggesting reduced digging behavior, which may indicate motor function deficit and/or behavioral changes. Mean ± SEM (n=30 total; control: n=18 (9 males, 9 females); mutant: n=12 (5 males, 7 females)). Unpaired *t*-test. **, p < 0.01. Created with BioRender.com. **E.** Open field test to measure locomotor activity and exploratory behavior. *Prx1-Rbm8a^f/+^*mice and control mice had comparable total distance traveled, with no significant difference observed. *Prx1-Rbm8a^f/+^*mice showed significantly reduced cumulative immobility time and significantly increased cumulative time spent in the border compared to controls. Representative EthoVision-generated movement tracks. Mean ± SEM (n=30 total; control: n=18 (9 males, 9 females); mutant: n=12 (5 males, 7 females)). Unpaired *t*-test. ns, not significant, *, p < 0.05. Created with BioRender.com. **F.** Elevated plus maze test to measure anxiety-like behavior and locomotor activity. *Prx1-Rbm8a^f/+^* mice and control mice had comparable total distance traveled, with no significant difference observed. *Prx1-Rbm8a^f/+^*mice showed significantly reduced cumulative immobility time compared to controls. *Prx1-Rbm8a^f/+^* mice and control mice showed no statistically significant difference in time spent in open arms. Representative EthoVision-generated movement tracks. Mean ± SEM (n=30 total; control: n=18 (9 males, 9 females); mutant: n=12 (5 males, 7 females)). Unpaired *t*-test. ns, not significant, *, p < 0.05. Created with BioRender.com. **G.** Representative images of P15 *Prx1-Rbm8a^f/+^* (right) and control (left) mouse brains. The brains of *Prx1-Rbm8a^f/+^* mice are noticeably smaller and differ in shape from those of littermate controls at P15. **H.** Quantification and statistical analysis of P15 brain weights of *Prx1-Rbm8a^f/+^* and control. Mean ± SEM (control: n=6; mutant: n=4). Unpaired *t*-test. **, p < 0.01. **I.** Representative coronal whole-brain MRI slices and fractional anisotropy maps from P15 control and *Prx1-Rbm8a^f/+^*mice. **J.** Quantification and statistical analyses of whole-brain volume and cortical volume from MRI analysis of P15 control and *Prx1-Rbm8a^f/+^* mice. Mean ± SEM (control: n=5; mutant: n=5). Unpaired *t*-test. ***, p < 0.001, ****, p < 0.0001.

Interestingly, in the nesting test, *Prx1-Rbm8a^f/+^* mice had significantly lower nesting scores at earlier timepoints but eventually caught up by later timepoints, indicating delayed nest-building behavior compared to controls^54^ (Supplementary Fig. 3D). Results from the forced swimming test^55^ and cylinder test^56,57^ also did not show significant differences between control and *Prx1-Rbm8a^f/+^* mice (Supplementary Fig. 3C, 3E). These results indicate that there are potential differences in the brains of *Prx1-Rbm8a^f/+^* mice that affect anxiety levels in novel environments (open field^52^ and elevated plus maze^58^) while still having normal coping behavior to acute stress (forced swimming^59^).

We next investigated whether these potential altered psychiatric states are consequences of changes in the brain structure and function. Given the significantly altered cranial development in *Prx1-Rbm8a^f/+^*mice (Figure 3I-3K), we hypothesized that the brains growing inside the cranium could be restricted and compressed, as in the case of craniosynostosis^60,61^.

Remarkably, the brains of *Prx1-Rbm8a^f/+^* mice were indeed noticeably smaller than those of littermate controls at P15 (Figure 4G). They were distinctly different in size and shape and significantly lighter than controls (Figure 4H). Magnetic resonance imaging (MRI) analysis confirmed statistically reduced whole-brain and cortical volumes in *Prx1-Rbm8a^f/+^* mice (Figure 4I, 4J). Taken together, these findings suggest that the severe skeletal defects in *Prx1-Rbm8a^f/+^* mice not only affect motor function but also brain shape and size, thereby potentially altering neural function.

### RBM8A-associated transcripts are significantly enriched in ciliary processes and developmental signaling pathways

RBM8A is an RNA-binding protein regulating multiple steps of mRNA processing, including mRNA splicing, export, degradation, and translation^62^. However, tissue-specific mRNAs that are bound to the EJC have yet to be identified. To investigate the downstream pathways controlled by RBM8A, we performed RIP-seq^63,64^ using embryonic mouse limb tissue to identify RBM8A-bound transcripts and to assess how that binding relates to our observed skeletal phenotype.

We used an anti-RBM8A antibody to immunoprecipitate protein-RNA complexes from E15 mouse forelimbs from wild-type mice, and isolated RNA from the co-precipitation for high-throughput sequencing. Our RIP-seq dataset identified a subset of transcripts with significant enrichment in RBM8A IP relative to input controls. A volcano plot of differential enrichment identified significantly enriched mRNAs linked to developmental regulators (Figure 5A). To elucidate the biological functions associated with the RBM8A-associated transcripts, we performed Gene Ontology (GO) Biological Process enrichment analysis^65^ on the significantly enriched genes. Enriched GO terms revealed strong representation of pathways related to cell cycle process, cilium organization and assembly, and neural tube development (Figure 5B).

**Figure 5:**
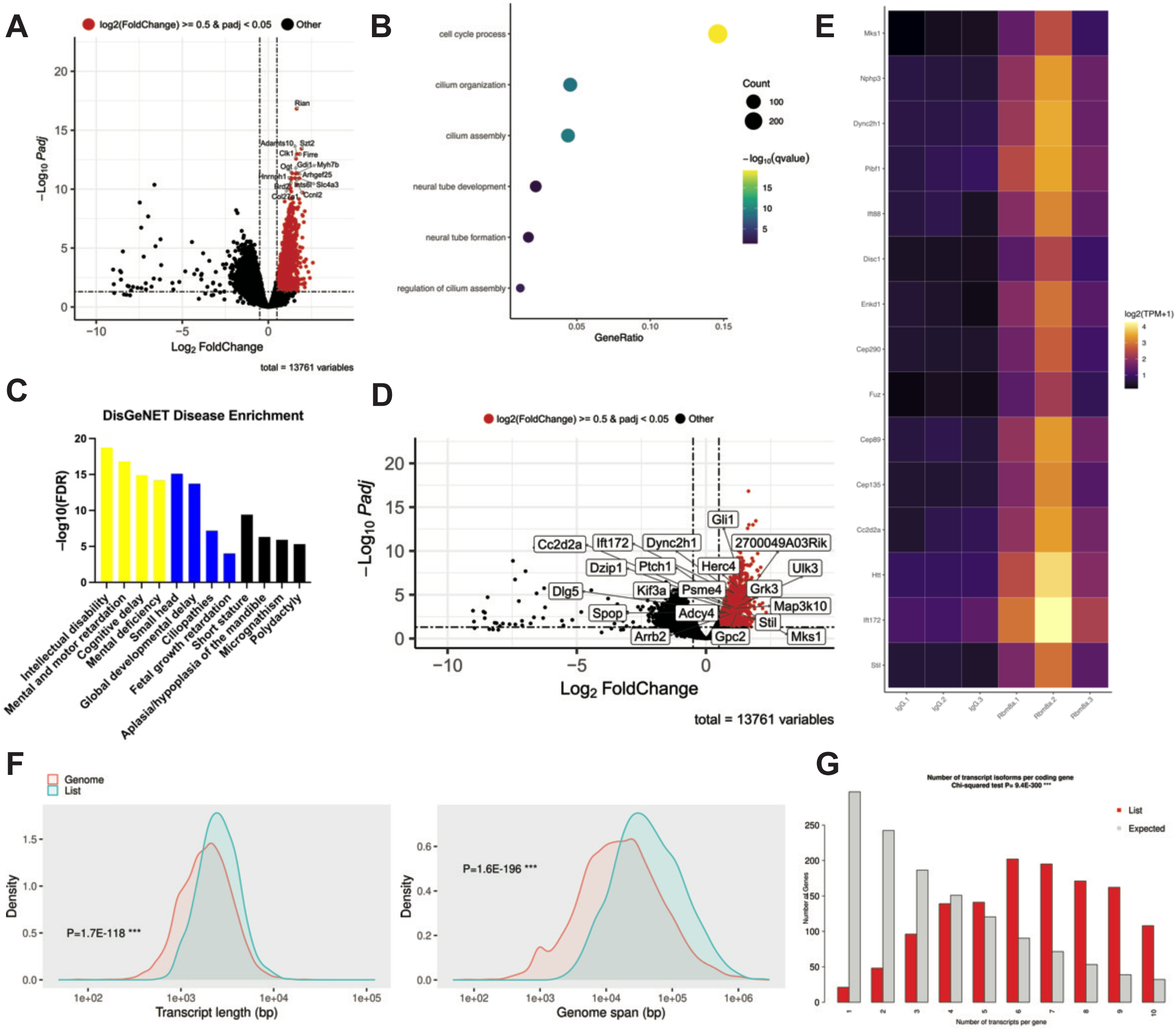
RIP-seq reveals enrichment of RBM8A-associated transcripts in ciliary processes and developmental signaling pathways. **A.** Volcano plot showing differentially enriched transcripts associated with RBM8A, identified by RIP-seq. The x-axis indicates log₂ fold change, and the y-axis represents −log₁₀ adjusted P value (q value). Significantly enriched transcripts are highlighted based on adjusted P value and fold-change thresholds. Transcripts meeting significance thresholds (adjusted *P* < 0.05 and log_2_|FC| ≥ 0.5) are highlighted in red. The gene names of the top 15 enriched transcripts are labeled. **B.** Selected GO biological process enrichment analysis of transcripts enriched in RBM8A immunoprecipitates. Enriched terms include cell cycle process, cilium organization, cilium assembly, neural tube development, neural tube formation, and regulation of cilium assembly. **C.** DisGeNET disease enrichment analysis of RIP-seq–identified transcripts enriched in RBM8A immunoprecipitates. Enriched disease terms were identified based on adjusted *P* value. Bars are color-coded to highlight major functional groupings: cognitive disability (yellow), general developmental conditions (blue), and craniofacial/skeletal abnormalities (black). **D.** Volcano plot showing differentially enriched transcripts associated with RBM8A, highlighting top 20 enrichment of transcripts involved in ciliary function and Hedgehog signaling pathways (e.g.,Ift172, Dync2h1, Ptch1, Gli1, Kif3a, and Mks1) from GO biological process, KEGG and Reactome databases. **E.** Heat map of transcripts enriched in RBM8A immunoprecipitates associated with primary cilia function showing relative expression across samples. Values represent normalized expressions and are displayed as row-wise z-scores. 3 replicates of IgG (control) on the left and 3 replicates from *Rbm8a* immunoprecipitates on the right. **F.** Transcript feature distribution plots show that RBM8A-associated transcripts (“list”; blue) were significantly enriched for longer transcript lengths and larger genomic span compared to background (“genome”; red). Chi-squared test. **G.** Distribution of the number of transcript isoforms per protein-coding gene for RBM8A-associated transcripts (“list”; red) compared to the genomic background (“expected”; gray). RBM8A-associated genes were significantly enriched for higher isoform counts, indicating a preference for genes with increased transcript complexity. Chi-squared test (P = 9.4 × 10⁻³⁰⁰).

Because these processes are associated with developmental and skeletal pathways, we next explored whether RBM8A-associated transcripts were enriched for genes implicated in human disease pathways using DisGeNET^66^. Notably, the top-most enriched disease terms were related to intellectual disability, general developmental conditions, and craniofacial/skeletal abnormalities (Figure 5C). Specifically, microcephaly, polydactyly, and craniofacial abnormalities are of meaningful relevance to TAR syndrome^19^. Additionally, enrichment of gene sets associated with ciliopathies is notable, as it is consistent with GO biological process enrichment for cilium organization and assembly (Figure 5B). This is further supported by the enrichment of genes involved in ciliary function and Hedgehog signaling (Figure 5D, 5E), suggesting that RBM8A-associated transcripts overlap with pathways regulating primary cilia function. To assess the consistency of pathway enrichment across multiple databases, we compared gene sets identified from GO, KEGG, and Reactome analyses, which revealed a shared subset of enriched transcripts of RBM8A in Hedgehog signaling pathways across all three databases (Supplementary Fig. 4F).

Following the identification of RBM8A-associated transcripts by RIP-seq, we next asked whether these transcripts exhibit distinct structural or genomic features that may explain their preferential binding with RBM8A. Transcript feature distribution plots show that RBM8A-associated transcripts were significantly enriched for longer transcript lengths compared to the background (Figure 5F). Consistent with this, RBM8A-associated transcripts were significantly enriched for genes with larger genomic spans, indicating that RBM8A preferentially binds to structurally complex, intron-rich genes (Figure 5F). Interestingly, RBM8A-associated transcripts show significant enrichment in a narrower and shorter 3’ UTR length range, suggesting that RBM8A binding is not driven by extended post-transcriptional regulatory regions (Supplementary Fig. 4A). Additionally, RBM8A-associated transcripts exhibited a modest but significant shift toward higher GC content, suggesting a mild sequence composition bias in those transcripts (Supplementary Fig. 4A). There was no significant difference in 5’ UTR length between RBM8A-associated transcripts and background, suggesting that RBM8A does not preferentially bind based on 5’ UTR structures (Supplementary Fig. 4A).

RBM8A-associated transcripts were strongly enriched for protein-coding genes, which is consistent with its established role in mRNA processing^23^ (Supplementary Fig. 4B). Consistently, these transcripts were significantly enriched for genes with more exons and depleted in genes with fewer exons, indicating preferential binding to transcripts requiring extensive splicing and its role in NMD^67–71^ (Supplementary Fig. 4C). Furthermore, these transcripts were significantly enriched for genes with more transcript isoforms, indicating preferential binding to structurally complex transcripts (Figure 5G). The chromosomal distribution plot shows RBM8A-associated transcripts had a broad distribution without chromosomal bias, suggesting that binding is based on RNA features rather than position (Supplementary Fig. 4D).

Based on our finding that RBM8A-associated transcripts exhibit increased structural complexity, we next examined whether these genes were enriched for RNA processing pathways. As expected, heatmap analysis revealed enrichment of genes involved in RNA splicing and processing (Supplementary Fig. 4E), suggesting that *Rbm8a* preferentially regulates structurally complex transcripts within ciliary and developmental gene networks through RNA processing mechanisms.

### *Rbm8a* deficiency alters cell fate and Hedgehog signaling in developing limbs

Given that RIP-seq analysis revealed enrichment of RBM8A-associated transcripts involved in cell cycle and ciliary function, we next asked whether disruption of *Rbm8a* causes early cellular defects relevant to limb development. To address this, we first performed immunocytochemistry on primary limb bud cells from *Prx1; Ai14^f/+^* mice (control) and *Prx1-Rbm8a^f/+^; Ai14^f/+^* mice (Figure 6A). Compared to control cells, we observed reduced fluorescence intensity of Ki67 (Figure 6B) and of Fibronectin (Figure 6C) in *Prx1-Rbm8a^f/+^; Ai14^f/+^* cells, suggesting altered proliferation and extracellular matrix organization^72–74^.

**Figure 6:**
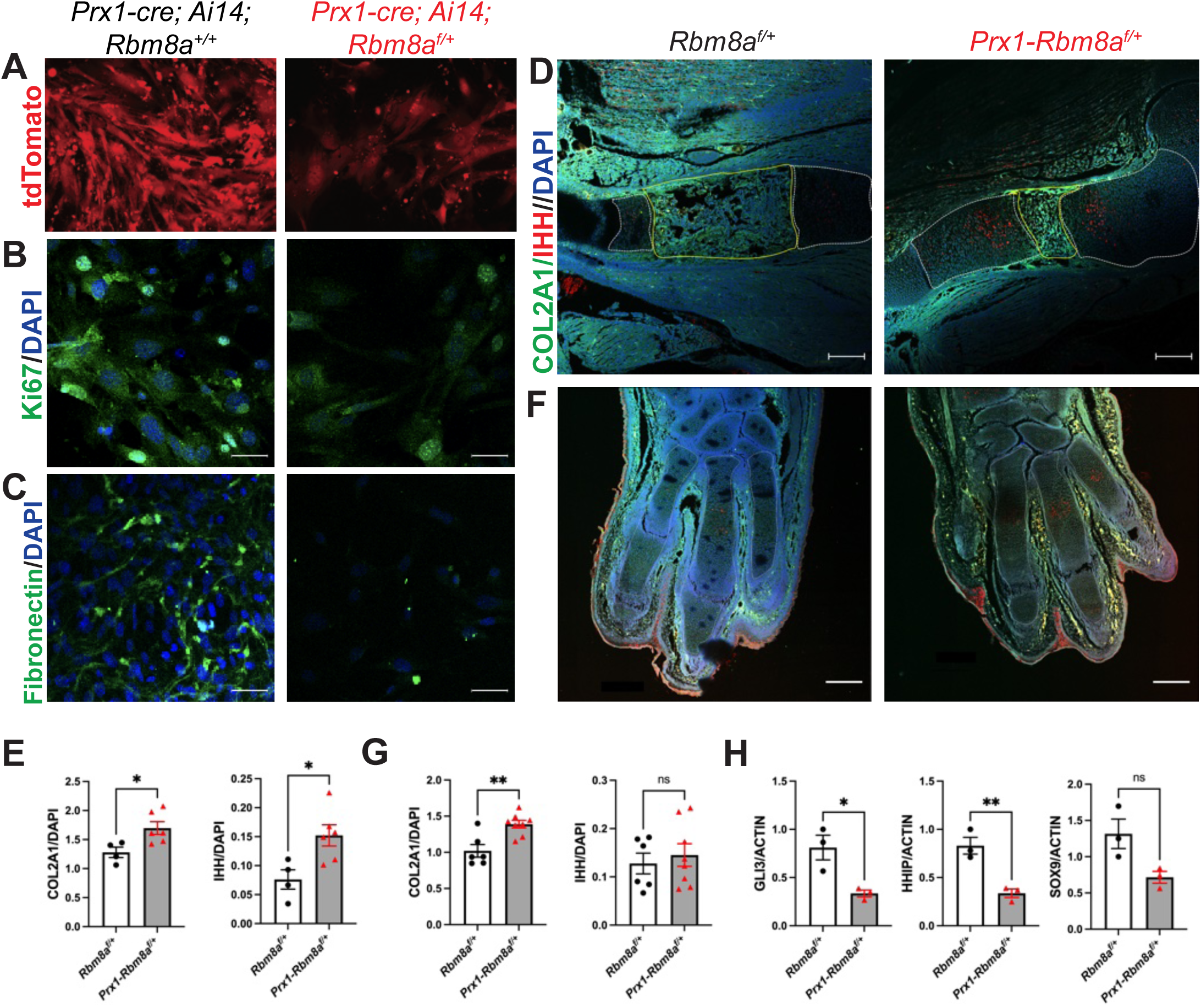
*Prx1-Rbm8a^f/+^* mice display disrupted ossification and Hedgehog signaling during embryonic limb development. **A.** Representative images of primary limb bud culture of *Prx1; Ai14^f/+^* mice (control) and *Prx1-Rbm8a^f/+^; Ai14^f/+^* mice showing Ai14 (tdTomato) reporter signal. **B.** Representative immunofluorescence staining images of primary limb bud culture of *Prx1; Ai14^f/+^*and *Prx1-Rbm8a^f/+^; Ai14^f/+^* mice showing Ki67 (green) merged with DAPI. Ki67 signal intensity is significantly reduced in *Prx1-Rbm8a^f/+^; Ai14^f/+^* cells compared to control cells. Scale bar = 50 µm. **C.** Representative immunofluorescence staining images of primary limb bud culture of *Prx1; Ai14^f/+^*and *Prx1-Rbm8a^f/+^; Ai14^f/+^* mice showing Fibronectin (green) merged with DAPI. Fibronectin signal intensity is significantly reduced in *Prx1-Rbm8a^f/+^; Ai14^f/+^* cells compared to control cells. Scale bar = 50 µm. **D.** Representative immunofluorescence staining images of E15 stylopods (humeri) from *Rbm8a^f/+^* and *Prx1-Rbm8a^f/+^* embryos showing COL2A1 (green), IHH (red), merged with DAPI (blue). The primary ossification center is outlined in yellow. Scale bar = 200 µm. **E.** Quantification and statistical analysis of COL2A1 and IHH signal intensity of control and *Prx1-Rbm8a^f/+^* stylopods. Both COL2A1 and IHH signal intensities are significantly higher in *Prx1-Rbm8a^f/+^* stylopods compared to those of controls. Mean ± SEM (control: n=4; mutant: n=6). Unpaired *t*-test. *, p < 0.05. **F.** Representative immunofluorescence staining images of E15 autopods (forepaws) from *Rbm8a^f/+^* and *Prx1-Rbm8a^f/+^* embryos showing COL2A1 (green), IHH (red), merged with DAPI (blue). Scale bar = 200 µm. **G.** Quantification and statistical analysis of COL2A1 and IHH signal intensity of control and *Prx1-Rbm8a^f/+^*autopods. COL2A1 signal intensity is significantly higher in *Prx1-Rbm8a^f/+^*autopods compared to those of controls. IHH signal intensity shows no significant difference between control and *Prx1-Rbm8a^f/+^* autopods. Mean ± SEM (control: n=6; mutant: n=8). Unpaired *t*-test. ns, not significant, **, p < 0.01. **H.** qRT-PCR results of Hedgehog pathway targets using RNA isolated from E15 forelimbs of *Prx1-Rbm8a^f/+^*and control embryos. Relative gene expression was quantified using the ΔΔCt method and normalized to Actin. *Gli3* and *Hhip* expressions were significantly reduced in *Prx1-Rbm8a^f/+^*relative to controls, with a similar trend observed in *Sox9* expression. Mean ± SEM (n=3 per group). Unpaired *t*-test. ns, not significant, *, p < 0.05, **, p < 0.01.

To examine whether these cellular changes were associated with disrupted chondrogenic progression and developmental signaling, we performed immunohistochemistry on E15 forelimbs from *Rbm8a^f/+^* (control) and *Prx1-Rbm8a^f/+^* mice. Both COL2A1 and IHH expression were significantly higher in *Prx1-Rbm8a^f/+^*stylopods, although COL2A1-positive domain appeared spatially altered, suggesting disrupted chondrocyte differentiation^75,76^ and a delayed or inefficient ossification center formation^9,77,78^ (Figure 6D, 6E). Additionally, *Prx1-Rbm8a^f/+^* mice had significantly higher COL2A1 expression (but not IHH) in the digits than control mice (Figure 6F, 6G). These results indicate disrupted early ossification processes and altered Hedgehog signaling in *Prx1-Rbm8a^f/+^*mice^12,79,80^.

To validate these findings at the mRNA level, we performed qRT-PCR analysis of Hedgehog pathway targets using RNA isolated from E15 forelimbs of *Prx1-Rbm8a^f/+^* and control mice. qRT-PCR revealed a significant reduction in *Gli3* and *Hhip* in *Prx1-Rbm8a^f/+^*forelimbs, indicating reduced downstream Hedgehog signaling activity^81–83^ (Figure 6H). Moreover, genes associated with chondrogenic and osteogenic differentiation, such as *Sox9* (Figure 6H), *Grem1*, and *Runx2*, showed downward trends in *Prx1-Rbm8a^f/+^*forelimbs (Supplementary Fig. 4G). In contrast, expression levels of *Foxf1* and *Ccnd1* were not significantly different between control and *Prx1-Rbm8a^f/+^*forelimbs (Supplementary Fig. 4G), suggesting that *Rbm8a* deficiency selectively affects specific components of developmental signaling pathways. Together, these findings suggest that *Rbm8a* deficiency disrupts Hedgehog-dependent cell differentiation during early limb development^74,84,85^.

### *Rbm8a* plays a role in additional organogenesis and hematopoiesis

Alterations in ciliary processes and developmental signaling pathways observed in our bone-specific cKO mouse model provide potential mechanistic insights into the absence of radius (skeletal defects) observed in TAR. Primary cilia are also present on almost all hematopoietic cells and are critical for regulating key signaling pathways for their proper development. Therefore, RBM8A-related defects in primary cilia and associated signaling pathways may also contribute to the platelet abnormalities observed in TAR.

Although *Prx1-Cre* drives recombination in limb and craniofacial mesenchyme (Figure 2A; Supplementary Fig. 1A), whether the defects in mesenchymal stromal cells in the bone marrow can cause indirect hematologic impairments is unknown. We examined whether thrombocytopenia is present in our *Prx1-Rbm8a^f/+^*mouse model. *Prx1-Rbm8a^f/+^* mice show no such effects on the complete blood count (CBC) test; their red blood cells, white blood cells, platelets, lymphocytes, neutrophils, and monocytes do not differ statistically from those of their littermate controls (Supplementary Fig. 5A). Their liver, kidney, and spleen are similar in size to those of their littermate controls, although their heart may be slightly smaller (Supplementary Fig. 5B).

To examine how mutations in *RBM8A* lead to platelet deficiency, we crossed *Pf4-Cre* mice with *Rbm8a^f/f^* mice to selectively delete *Rbm8a* in megakaryocytes and platelets (Supplementary Fig. 6A). Interestingly, homozygous *Pf4-Rbm8a^f/f^* embryos were viable and exhibited normal survival and development into adulthood. As expected, *Pf4-Rbm8a^f/+^*mice exhibit a significant reduction in platelet counts, with no differences in white blood cell, lymphocyte, neutrophil, or monocyte counts (Supplementary Fig. 6B). Strikingly, *Pf4-Rbm8a^f/f^* mice showed an even more severe reduction in platelet counts (Supplementary Fig. 6B). The liver, kidney, and heart of *Pf4-Rbm8a^f/+^* mice were comparable to those of their littermate controls. Intriguingly, the spleen of homozygous *Pf4-Rbm8a^f/f^*mice had noticeably enlarged spleens (splenomegaly) compared to those of their littermate controls and heterozygous *Pf4-Rbm8a^f/+^* mice (Supplementary Fig. 6C). This result further supports previous studies that RBM8A is required for normal hematopoietic processes^86,87^. Further studies utilizing this blood-specific cKO mouse model can elucidate the role of *Rbm8a* in hematopoiesis.

## Discussion

Here, we identify a novel role for *Rbm8a* in regulating skeletal development through its role in RNA processing of structurally complex transcripts. Because *RBM8A* mutations cause TAR syndrome^22^, we developed a mouse model to investigate how *Rbm8a* deficiency results in the TAR skeletal phenotype. We validate the first mouse model of TAR syndrome, *Prx1-Rbm8a^f/+^*, by various phenotypic, behavioral, and molecular analyses. Our findings reveal new insights into structural and functional changes in these mouse brains, which are linked to neurological phenotypes reported in human TAR patients. Our RIP-seq analysis revealed that *Rbm8a* preferentially binds with long, exon-rich transcripts with more isoforms, which are enriched in ciliary and developmental signaling pathways. Immunostaining and qRT-PCR results showed the selective downregulation of Hedgehog target genes in our mutant mice. Together, our findings support a model in which *Rbm8a* regulates RNA processing of transcripts that are enriched in primary cilia and developmental signaling pathways, and pinpoints the disruption of this process as the mechanism by which *Rbm8a* deficiency leads to impaired Hedgehog signaling, contributing to skeletal maldevelopment in TAR syndrome.

### Role of *Rbm8a* in primary cilia and Hedgehog signaling function

Primary cilia are non-motile sensory organelles that function as cellular antennae to sense and transduce extracellular signals and regulate multiple core signaling pathways^88,89^. During osteogenesis, primary cilia are essential in coordinating osteoblast differentiation, chondrocyte proliferation, and bone matrix formation^88,90^. Because primary cilia consist of many interdependent protein complexes that are tightly assembled, they are vulnerable to transcriptional changes^88^. More specifically, many ciliary genes have long introns, many exons, and complex splicing patterns, rendering them particularly sensitive to RNA processing defects^91,92^. Given that *RBM8A* controls post-transcriptional RNA processing in its role in the EJC^23^, *RBM8A* deficiency likely results in primary ciliary signaling defects (Figure 5; Supplementary Fig. 4). These perturbations subsequently dysregulate core developmental signaling pathways (such as Hedgehog), leading to phenotypic alterations, as observed in this study. The skeletal phenotypes of *Prx1-Rbm8a^f/+^* mice align with those observed in human skeletal ciliopathies such as Jeune syndrome, Ellis–van Creveld syndrome, cranioectodermal dysplasia, and short-rib polydactyly syndromes^93,94^, further suggesting that disrupted primary cilia-dependent signaling pathways result in congenital bone defects.

As such, our findings indicate that *Rbm8a* deficiency alters the expression of genes associated with the Hedgehog signaling pathway. Significantly reduced *Gli3* and *Hhip* expression (Figure 6H) in *Prx1-Rbm8a^f/+^* indicate weakened Hedgehog signaling^80–83^. Downward trends, but not statistically significant attenuation of, downstream regulators such as *Sox9*, *Grem1*, and *Runx2*^74,84,85^ (Figure 6H; Supplementary Fig. 4G), suggest that the suppression of pathway activity is moderate. Surprisingly, increased IHH expression was observed in the pre-hypertrophic/hypertrophic chondrocyte region next to the primary ossification center in the humerus (Figure 6D, 6E). IHH may be upregulated as a compensatory response to attenuated Hedgehog pathway activity^9,95,96^. Another potential explanation is that *Prx1-Rbm8a^f/+^* mice produce more ligand (IHH) but respond less effectively (*Gli3* and *Hhip*)^95,97^. This would prevent proper progression of osteogenesis, resulting in reduced expansion of the primary ossification center (Figure 6D) and a delay in ossification^98^.

### Role of *Rbm8a* in regulation of osteogenesis

Osteogenesis has two main processes: endochondral and intramembranous^9,11^. Long bones, such as the limb bones, form through endochondral ossification, in which a cartilage template gets replaced by bone^9,11^. Most cranial bones develop through intramembranous ossification, in which mesenchymal cells directly differentiate into osteoblasts, though some cranial bones form via endochondral ossification^9,11^. Our study shows that in *Prx1-Rbm8a^f/+^* mice, there are significant alterations in long bones (digits, radii, ulnae, humeri, tibia, and femur) and various cranial landmarks (Figure 2; Figure 3; Supplementary Fig. 2). The fact that both ossification processes are disrupted by *Rbm8a* deficiency suggests that this gene is important in both mechanisms. Notably, both osteogenesis processes involve conserved developmental networks such as Hedgehog signaling^9,11,79^, demonstrating how disruption of shared molecular regulators can yield context-dependent, tissue-specific phenotypes. Furthermore, embryonic lethality observed in homozygous *Prx1-Rbm8a^f/f^*mice (Figure 2F) indicates a critical role of *Rbm8a* in developmental processes required for fetal viability in mice, consistent with the key role of primary cilia-dependent Hedgehog signaling across multiple organ systems during development^81,88^.

Notably, *Prx1-Rbm8a^f/+^* mice exhibit significant alterations in brain structure (Figure 4G-4J). This is likely due to constraints imposed by the significant cranial abnormalities, which are present as early as P5 and continue to at least P30 (Figure 2I-2S; Supplementary Fig. 2A-2F). This restriction would prevent the brain growing inside the cranium from developing properly, analogous to craniosynostosis^60,61^. Subsequent changes in brain size and shape likely affect its function, as reflected in some of the behavioral tests we performed (Figure 3E, 3F; Supplementary Fig. 3). This resembles the neurological findings reported in some TAR patients, such as intellectual disability and psychiatric disorders^19^. Due to the rarity of TAR syndrome, the neurological aspects of the condition are not fully defined. Our findings can serve as a foundation for further understanding of the distinct ways in which TAR brains may be altered. Moreover, microcephaly and intellectual disability were among the top-enriched disease terms for RBM8A-associated transcripts (Figure 5C). These conditions are also observed in 1q21.1 deletion syndrome^99^, highlighting that different deletions within the same chromosomal region yield overlapping phenotypes, though via distinct mechanisms.

### Role of RNA regulation in ossification

Beyond the context of *RBM8A* and TAR syndrome, our findings illuminate how perturbations in RNA regulatory mechanisms can affect a developmental signaling program that guides embryonic bone formation by governing signaling robustness. As such, this study can serve as a potential framework for understanding the mechanisms of other congenital skeletal disorders. Our study supports the core concept that developmental signaling pathways function in a tightly coordinated regulatory network that combines transcriptional and post-transcriptional inputs^23^. Accordingly, it is through the RNA quality control function of *RBM8A* that the stability of transcripts necessary for proper Hedgehog signaling responses is affected. As a consequence of altered RNA regulatory function and subsequent changes in transcript stability, cellular signaling competence may be impacted^23,100^. When cellular signaling competence is reduced, cells are less effective in properly interpreting and transducing signaling pathway cues^101^. This explains the moderate (not fully abolished) molecular changes and variable skeletal phenotypes observed in this study. Our findings highlight the modulation of RNA processing mechanisms as a critical contributor to proper ossification processes during early skeletal development.

While this study provides novel insights into *Rbm8a* and bone development, some limitations should be considered. While our RIP-seq analysis identified RBM8A-associated transcripts, it cannot define individual targets that directly impact the observed skeletal phenotypes. Our qPCR analyses of several Hedgehog-related genes confirmed the attenuated Hedgehog pathway, but testing additional genes in the pathway is necessary to understand the broader scope. Future studies should include bulk RNA-sequencing comparing control and *Prx1-Rbm8a^f/+^* mouse limbs to investigate overall transcriptional alterations in relevant genes, thereby bridging RIP-seq and qPCR data. Moreover, examining the impact of *Rbm8a* deficiency in other primary cilia-dependent pathways involved in osteogenesis (Wnt, TGF-β/BMP, and PDGF)^88,90^ can help determine the extent of primary ciliary signaling defects. Additionally, single-cell RNA-sequencing would help reveal potential cell-type-specific perturbations in the altered Hedgehog signaling pathway. Lastly, additional experiments to parse out the neurological aspects of *Prx1-Rbm8a^f/+^*mice can demonstrate a new understanding of TAR brains. Behavioral tests for learning and memory can reveal novel insights into how the distinct brain changes affect cognition, and examining cellular changes in cranial bones can help explain those behavioral outcomes.

In all, our present study establishes *Rbm8a* as a critical regulator of embryonic bone development by maintaining the integrity of structurally complex transcripts necessary for primary cilia and developmental signaling pathways. We demonstrate that disruption in RNA processing (i.e., *Rbm8a* deficiency) alters Hedgehog signaling, leading to significantly impaired ossification processes and subsequent skeletal maldevelopment. These findings provide new insight into the mechanisms underlying not only TAR syndrome, but also other congenital skeletal anomalies and other *RBM8A*-related disorders. Taken together, our study provides a framework for future research examining the impact of RNA processing pathways during early development, and for translational studies addressing human congenital skeletal disorders.

## Methods

### Human Bone Marrow-Derived MSC Culture

Human bone marrow-derived MSCs were obtained from ATCC (catalog #: PCS-500-012) and cultured using Bone Marrow-Mesenchymal Stem Cell Basal Medium (ATCC, catalog #: PCS-500-030) with 7% FBS and growth factor supplements (15 ng/ml IGF-1, 125 ng/ml FGF2).

### Generation of the *Prx1*-*Cre*; *Rbm8a^f/+^* Mice

*Rbm8a*-floxed mice (*Rbm8a^f/f^*) without the Neo cassette were previously generated in our lab^25,26,28^. These mice were then crossed with *Prx1-Cre* (Jackson Laboratory, stock #: 005584) to selectively knock out *Rbm8a* in early limb bud mesenchyme and a subset of craniofacial mesenchyme. The resulting *Prx1-Cre; Rbm8a^f/+^* mice display significant bone development defects but are otherwise healthy. However, we cannot detect *Prx1-Cre; Rbm8a^f/f^* mice at the postnatal stage, as they are embryonic lethal around E12-13.

### Generation of the Pf4-Cre; Rbm8a^f/+^ and Pf4-Cre; Rbm8a^f/f^ Mice

*Rbm8a*-floxed mice (*Rbm8a^f/f^*) without the Neo cassette were previously generated in our lab^25,26,28^. These mice were then crossed with *Pf4-Cre* (Jackson Laboratory, stock #: 008535) to selectively knock out *Rbm8a* in megakaryocytes. The resulting *Pf4-Cre; Rbm8a^f/+^* and subsequent *Pf4-Cre; Rbm8a^f/f^* mice display significant reductions in platelet counts.

### Animals

All procedures involving mice were reviewed and approved by the Pennsylvania State University IACUC committee under IACUC protocols 44057 and 800335. Mice were housed by sex (2–5 mice per cage) in a room with a 12-hour light/dark cycle and provided *ad libitum* access to food and water.

### Timed Mating and Embryo Collection

Timed matings were performed by housing a *Prx1-Cre; Rbm8a^f/+^*male and a *Prx1-Cre; Rbm8a^f/+^* female mouse together overnight, then designating the presence of a vaginal plug the following morning as E0.5. Pregnant female mice were euthanized at E15, and the embryos were harvested in cold PBS. Both forelimbs of each embryo were carefully isolated using forceps and a microscope, then stored immediately in -80 °C.

### Primary Limb Bud Cell Culture

*Prx1-Cre; Rbm8a^f/+^* mice were crossed with Ai14 reporter mice (Jackson Laboratory, stock #: 007914) for the desired genotypes of *Prx1-Cre*; Ai14*^f/+^* mice (control) and *Prx1-Cre; Rbm8a^f/+^*; Ai14*^f/+^* mice. Pregnant female mice were euthanized at E11-E12, and the embryos were harvested in a sterile Petri dish with ice-cold PBS. Both limb buds of each embryo were carefully isolated using forceps and a microscope, then transferred to a new dish with sterile PBS. The limb buds were minced using fine scissors, then incubated in 0.25% trypsin at 37°C for 15 minutes. The samples were further dissociated by gently pipetting up and down. The digestion was stopped by adding an equal volume of Gibco DMEM/F-12 medium (Thermo Fisher Scientific, catalog #: 11320033), supplemented with 10% FBS and 1% Penicillin-Streptomycin. This cell suspension mixture was passed through a 70 μm cell strainer to remove debris, then plated directly onto gelatin-coated glass coverslips. The cells were incubated at 37°C and 5% CO₂, with media changes every 2-3 days.

### Immunocytochemistry

Sterile glass coverslips were incubated in EmbryoMax 0.1% gelatin solution (MilliporeSigma, catalog #: ES-006-B) for 2 hours at 37°C and 5% CO₂, then washed 3 times with sterile PBS and air-dried in the tissue culture hood. After the primary limb bud cells were plated and grown on these coverslips, the staining was performed as previously described^25^. Briefly, they were fixed with 4% PFA for 15 minutes at room temperature, then permeabilized by incubation in 0.3% Triton X-100 solution for 1 hour at room temperature. Subsequently, they were incubated in 5% normal serum for 1 hour at room temperature, then incubated overnight at 4 °C in rabbit anti-RBM8A (GeneTex, catalog #: GTX131387), mouse anti-Fibronectin (DSHB Hybridoma, catalog #: P1H11), mouse anti-N-Cadherin (DSHB Hybridoma, catalog #: MNCD2-s), rabbit anti-Ki67 (Invitrogen, catalog #: 701198), or goat anti-SOX9 (R&D Systems, catalog #: AF3075). After staining, coverslips were mounted onto microscope slides using ProLong Diamond Antifade Mountant with DAPI (Invitrogen, catalog #: P36971). Whole slide scanning (20x) was performed on the Zeiss LSM 800 Confocal Laser Scanning Microscope.

### Immunohistochemistry

Wild-type E10.5 and E14.5 mouse embryos were fixed in 4% PFA in PBS for 1 hour, briefly washed with PBS, then left in 30% sucrose overnight, embedded in O.C.T freezing media, and stored at −80 °C. Cryosections at 10 µm thickness were cut with a Leica LM1900 Cryostat. E15 *Prx1-Rbm8a^f/+^* and control embryos were collected from pregnant dams, and the forelimbs of those E15 mice were isolated using forceps and a microscope. The forelimbs were then postfixed for 24 hours in 4% PFA at 4 °C, then stored in PBS at 4 °C. Samples were processed and embedded in paraffin by HistoWiz Inc. (Long Island City, NY, USA) using their standard operating procedure. The paraffin-embedded samples were sectioned at 5 μm using the Thermo Shandon Finesse paraffin microtome, then dewaxed using the Leica Autostainer ST5010 XL at the Huck Institutes Microscopy Facility (University Park, PA, USA). The samples were then incubated in Pepsin-HCl (Mettler Toledo, catalog #: 01-912-587) for 30 minutes at 37 °C for antigen retrieval. The staining was performed as previously described. Antibodies used include: CD-90 (Invitrogen, catalog #: 14-0909-82), rabbit anti-RBM8A (GeneTex, catalog #: GTX131387), goat anti-SOX9 (R&D Systems, catalog #: AF3075), rabbit anti-IHH (Proteintech, catalog #: 13388-1-AP), mouse anti-COL2A1 (DSHB Hybridoma, catalog #: II-II6B3). After staining, sections were coverslipped using ProLong Diamond Antifade Mountant with DAPI (Invitrogen, catalog #: P36971). Whole slide scanning (20x) was performed on the Zeiss LSM 800 Confocal Laser Scanning Microscope.

### Behavioral Tests

All behavioral tests were performed in a sound-attenuated room. A cohort of 30 mice that are similar in age was used. They were 2-3 months old on the first behavioral test and 4-5 months old at the last test. The cohort consisted of the following: *Rbm8a^f/+^*: n=9 females, n=9 males; *Prx1-Cre; Rbm8a^f/+^*: n=7 females, n=5 males.

### Open Field Test

Mice were placed in the center of the open field arena (40 cm x 40 cm) and allowed to move freely for 10 minutes while being tracked by the Noldus EthoVision XT video behavior recognition system. At the conclusion of each trial, the surface of the arena was cleaned with ethanol.

### Elevated Plus Maze

Mice were placed in a four-arm maze that is elevated 50 cm above the floor. In this apparatus, the four arms intersect to form a plus shape. Two of the arms are closed on the sides, and the other two arms are open. Mice were placed in the center of the apparatus, facing an open arm, and allowed to freely explore the maze for 10 minutes while being tracked by the Noldus EthoVision XT video behavior recognition system. At the conclusion of each trial, the surface of the arena was cleaned with ethanol.

### Rotarod

Mice were placed on a rotating drum with controlled, accelerating speed (4-40 rpm over 300 seconds). When the mice fall approximately 8 inches from the rotating drum onto the lever, their body weight presses the lever down, automatically pausing the test timer. The time each mouse took to fall was recorded. Each mouse was tested three times per day, and the latency to fall was averaged from those trials. All mice are tested for three consecutive days to test motor learning and memory.

### Cylinder Test

Mice were placed in a clear glass cylinder (8 cm in diameter and 11 cm in height) and video-recorded for 10 minutes. The videos were manually scored by three lab members (blinded to genotype) for independent, weight-bearing contacts on the side of the cylinder wall to assess forelimb asymmetry. The results were averaged, and percentages of left and right paw contact, as well as both paw contact, were calculated.

### Forced Swimming

Mice were individually placed in a transparent cylinder (8 cm in diameter and 11 cm in height) filled with water (25 ± 1°C) while being video-recorded. Each session lasted for 6 minutes, and the duration of immobility was recorded during the final 4 minutes. Immobility was defined as the absence of active escape behaviors, with only minimal movements necessary to keep its head above water. Time spent being immobile was manually scored using the video recording by three separate lab members blinded to genotype, and the results were averaged. Water was changed between mice, and the water temperature was checked with a thermometer before each trial.

### Marble Burying Test

Each mouse was placed in a new cage with 3 cm of bedding material with five rows of four marbles on the surface of the bedding. The mice were allowed to move freely in the cage for 10 minutes. At the conclusion of each trial, the cages were photographed to document the status of the marbles. Three separate lab members blinded to genotype independently scored the number of marbles buried, and the results were averaged.

### Grip Strength

Mice were handled by the tail and allowed to grasp the test bar with their forepaws, then gently pulled backward by the tail in a horizontal plane until they released the grip. The maximum force exerted before release was recorded using a grip strength meter. Each mouse underwent six consecutive trials with 1-minute rests between trials to minimize fatigue. Grip strength was normalized by the weight of each mouse, and the average of six trials was used for analysis.

### Grid Walking Test

Mice were placed on a smooth, rounded wire grid (2 mm wire diameter) that was slightly elevated (10 cm) from the bottom of the apparatus. The mice were allowed to freely explore the arena for 5 minutes while being video-recorded from below the grid wire surface. Three separate lab members blinded to genotype independently counted the number of paw slips (paw fully going through the grid), and the results were averaged.

### Nesting Test

Mice were individually housed in clean cages and provided with a single pre-weighed compressed cotton nesting material. Nesting was evaluated at 1, 3, 6, 24, and 36 hours following the introduction of nesting material, with a photo record at each time point. The scoring was performed using those photos with a standardized 5-point scale as previously described^54^. Three separate lab members blinded to genotype independently evaluated the nesting score, and the results were averaged.

### Micro-computed Tomography (micro-CT)

Postnatal mice were imaged *ex vivo* using SkyScan 1176 (Bruker, Billerica, MA, USA) in the Huck Institutes High Field Magnetic Resonance Imaging Core Facility (University Park, PA, USA). The micro-CT projection images were reconstructed into cross-section images using NRecon (Bruker, Billerica, MA, USA). The cross-section images were then analyzed using Avizo (Thermo Fisher Scientific, Waltham, MA, USA).

### Complete Blood Count (CBC) Test

A minimum of 40 uL blood was collected from each adult mouse via cardiac puncture under terminal anesthesia and collected into BD Microtainer blood collection tubes (BD, catalog #: 365974). The CBC test was performed using ProCyte Dx Veterinary CBC Hematology Analyzer (IDEXX, Westbrook, ME, USA) by the Penn State Animal Resource Program staff.

### RNA Isolation

E15 forelimb tissues were homogenized using a mini handheld plastic tube pestle in 1.5 mL microcentrifuge tubes. A set of forelimbs per embryo was used as a single sample. RNA was isolated using the RNeasy Mini Kit (QIAGEN, catalog #: 74104) according to the manufacturer’s protocol. The RNA yields were assessed using the NanoDrop microvolume spectrophotometer (Thermo Fisher Scientific, Waltham, MA, USA).

### Quantitative Reverse Transcription–PCR (qRT-PCR)

The total RNAs were converted into cDNA by reverse transcription with the oligo-dT primer, using Superscript reverse transcriptase III (Invitrogen, catalog #: 18080093). qRT-PCR was performed using PowerTrack SYBR Green Master Mix for qPCR (Applied Biosystems, catalog #: A46109) with the following primers (5′–3′): mouse RBM8A primers (F: ATTACGACAGTGTGGAGCAG, R1: TTGACCATTTAGTCCTTCCA), GLI3 (F: GGAGCAGGTTCTGGAACCTT, R: TCCATGTTCTGGGCAGTCTT), HHIP (F: TTGCCACTCAGGAGGTTGAA, R: TTCTGGTGCTCAGGACTGGT), SOX9 (F: CAGCAAGACTCTGGGCAAGC, R: GGTGGTCCTTTCCTCTTGTG), GREM1 (F: TCTTCTCCGTGTCCTCCTGA, R: GTGAGCGTAGTGAGCAGCAG), RUNX2 (F: GACTGTGGTTACCGTCATGGC, R: ACTTGGTTTTTCATAACAGCGGA), CCND1 (F: GCAAGTCAAGAGGAGCAGGA, R: GGAAGAGGTAGGTGGAGGAG). The expression of the β-Actin gene (F: CGTGGGCCGCCCTAGGCACCA, R: TTGGCCTTAGGGTTCAGG GGGG) was used as an internal control. Primers were designed using NCBI Primer-BLAST and synthesized by IDT. The qPCR reactions were performed on a StepOnePlus Real-Time PCR system (Applied Biosystems, Carlsbad, CA, USA), and the Ct calculations were performed using StepOne software version 2.3.

### RNA Immunoprecipitation

Roughly 100 E15 embryos were collected from multiple pregnant dams. Both forelimbs from each E15 embryo were isolated and pooled to yield 3 total replicates. The pooled samples were lysed in homogenization buffer at 2% weight/volume using a mini handheld plastic tube pestle in 1.5 mL microcentrifuge tubes. Each homogenate was centrifuged at 10,000 rpm for 10 minutes at 4°C. Supernatants were transferred to fresh tubes, and either rabbit anti-RBM8A antibody (GeneTex, catalog #: GTX131387) or anti-IgG antibody was added to each tube, which were incubated in the cold room for 4 hours with rotation. Dynabeads Protein G for Immunoprecipitation (Invitrogen, catalog #: 10004D) was equilibrated to the homogenization buffer using DynaMag-2 Magnet (Invitrogen, catalog #: 12-321-D). The antibody-supernatant mixtures were added to pre-cleared Dynabeads Protein G and incubated overnight at 4 °C with rotation. The next day, the antibody-supernatant-bead mixtures were placed on the DynaMag-2 Magnet to remove the supernatant. The beads were washed with a high-salt buffer three times, each for 10 minutes at 4 °C with rotation. After the last wash, lysis buffer (10 μL β-mercaptoethanol per mL RLT lysis buffer) was immediately added to the beads, and the mixture was vortexed for 30 seconds. The samples were then placed on the DynaMag-2 Magnet, and the supernatant was collected and transferred to new tubes. RNA was isolated from these samples using the RNeasy Mini Kit (QIAGEN, catalog #: 74104) according to the manufacturer’s protocol. The RNA yields were assessed using the NanoDrop microvolume spectrophotometer (Thermo Fisher Scientific, Waltham, MA, USA).

### RIP-seq – Library Preparation & Sequencing

RIP-seq library preparation and sequencing were performed by Azenta (Chelmsford, Massachusetts, USA). RNA samples were quantified using Qubit 4.0 Fluorometer (Thermo Fisher Scientific, Waltham, MA, USA), and RNA integrity was assessed using TapeStation (Agilent Technologies, Palo Alto, CA, USA). The NEBNext Ultra II RNA Library Prep Kit for Illumina (New England Biolabs, Ipswich, MA, USA), including clustering and sequencing reagents, was used for library preparation without RNA enrichment or depletion, according to the manufacturer’s recommendations. Briefly, total RNA was directly fragmented and primed using random primers. RNA fragmentation was performed at 94 °C for a duration selected based on RNA integrity to target an average insert size of approximately 200 nucleotides. First-strand and second-strand cDNA were subsequently synthesized. The resulting cDNA fragments were end-repaired and adenylated at 3’ ends, and universal adapters were ligated to the cDNA fragments, followed by index addition and library enrichment by PCR with a limited number of cycles. The sequencing library was validated on the Agilent TapeStation (Agilent Technologies, Palo Alto, CA, USA) and quantified using Qubit 4.0 Fluorometer (Thermo Fisher Scientific, Waltham, MA, USA) and qPCR (KAPA Biosystems, Wilmington, MA, USA).

The sequencing libraries were multiplexed and clustered onto a flowcell on the Illumina NovaSeq instrument according to the manufacturer’s instructions. The samples were sequenced using a 2x150bp Paired End (PE) configuration. Image analysis and base calling were conducted by the NovaSeq Control Software (NCS). Raw sequence data (.bcl files) generated from Illumina NovaSeq were converted into fastq files and de-multiplexed using Illumina bcl2fastq 2.20 software. One mismatch was allowed for index sequence identification.

### RIP-seq – Data Analysis

After assessing the quality of the raw data, sequence reads were trimmed to remove possible adapter sequences and nucleotides with poor quality. The trimmed reads were mapped to the reference genome available on ENSEMBL using the STAR aligner v.2.5.2b. The STAR aligner is a splice aligner that detects splice junctions and incorporates them to help align the entire read sequences. BAM files were generated from this step. Unique gene hit counts were calculated by using feature Counts from the Subread package v.1.5.2. Only unique reads that fell within exon regions were counted. After the extraction of gene hit counts, the gene hit counts table was used for downstream differential expression analysis. Using DESeq2, a comparison of gene expression between the sample groups was performed. The Wald test was used to generate p-values and log2 fold changes. Genes with adjusted p-values (Padj) <= 0.05 and log2 fold changes >= 0.5 were called as differentially expressed genes for each comparison.

Differential exon usage between experimental groups was analyzed using DEXSeq v. 1.46.1, and exon bins with adjusted p-values (Padj) <= 0.05 and log2 fold changes >= 0.5 were considered significantly differentially used. Gene ontology and pathway enrichment analyses were performed on significantly differentially expressed genes and significantly differentially used exons using clusterProfiler v.4.10.0 together with org.Mm.eg.db v.3.17.0. Enrichment was assessed across GO, KEGG, and Reactome databases. Volcano plots were generated using EnhancedVolcano v. 1.18.0. Disease enrichment analysis was performed using enrichR v. 3.4. Mouse gene symbols were used as input for enrichment analysis against the DisGeNET disease annotation library. All downstream analyses were performed in R 4.3.1.

### MRI - Acquisition

All samples were imaged using a 7T MRI system with a quadrature transmit volume coil (70 mm diameter) and a four-channel receive-only phased array cryocoil (Bruker Biospin, Billerica, MA, USA) at the Bernard and Irene Schwartz Center for Biomedical Imaging (New York, NY, USA). Co-registered T2-weighted images were acquired using a RARE sequence with the following parameters: TE/TR = 50/3000 ms, one signal average, echo train length = 8, the same FOV, matrix size, and slice thickness as the diffusion tensor images. Co-registered diffusion tensor images were acquired using a diffusion-weighted echo planar imaging (DW-EPI) sequence with the following parameters: TE/TR = 30/5000 ms, one signal average, four segments, diffusion gradient duration = 5 ms, diffusion time = 15 ms, 30 diffusion encoding directions, b = 1000, 2000 s/mm2, five non-diffusion-weighed images, FOV = 15 × 15 mm, matrix size = 128 × 128, slice thickness = 0.5 mm, and an in-plane resolution = 0.117 × 0.117 mm2.

### MRI - Volume Measurement

Whole brain and cortex volumes were determined using segmentation analysis (AMIRA 3D, 2024.1; Thermo Fisher Scientific) software. The brain and cortex were manually outlined and segmented in AMIRA. The number of voxels from segmented tissue within each slice was added together to compute the total volume (= num of voxels * volume unit of voxels) for each brain region and the total volume. The computation was carried out in MATLAB R2025a.

### Tissue Optical Clearing

Commercially available SHIELD (Stabilization under Harsh conditions via Intramolecular Epoxide Linkages to prevent Degradation) preservation, passive clearing reagents, and detailed protocols were obtained from LifeCanvas Technologies (Cambridge, MA, USA). Samples were incubated in SHIELD OFF solution for 3 days at 4 °C on an orbital shaker. Subsequently, the SHIELD OFF solution was replaced with SHIELD ON buffer and incubated for 24 hours at 37 °C with shaking. Tissues were incubated in 20 mL of delipidation buffer with 10 μl of SYTOX™ Deep Red Nucleic Acid Stain (Thermo Fisher Scientific, catalog #: S11381) at 37 °C for 7 days. To match the refractive index (RI) of the delipidated tissues (RI = 1.52), samples were incubated in 20 mL of 50% EasyIndex (LifeCanvas Technologies, catalog #: EI-Z1001) + 50% distilled water for 1 day, then switched to 100% EasyIndex solution for another day at 37 °C with gentle shaking.

### Lightsheet Fluorescence Microscopy Imaging

Tissue optical cleared samples were embedded in an agarose solution containing 2% low-melting agarose (MilliporeSigma, catalog #: A2576) in EasyIndex using a custom sample holder. Embedded samples were incubated in EasyIndex at 37 °C overnight, then imaged using the SmartSPIM lightsheet fluorescence microscope (LifeCanvas Technologies, Cambridge, MA, USA). During imaging, the sample holder arm supporting the embedded sample was immersed in an index-matched immersion liquid. The imaging setup consisted of a 3.6X objective lens (LifeCanvas Technologies, 0.2 NA, 12 mm working distance, 1.8 μm lateral resolution), three lasers at 488 nm, 561 nm, and 642 nm, and a 5 μm z-step size. Maximum intensity projections were generated for visualization. For quantification, Fiji (ImageJ) was used to define regions of interest (ROIs) for the forelimbs and crania, then the total tissue area and Ai14 signal-positive area were measured. The area-normalized Ai14 signal intensity was calculated as the ratio of Ai14 signal-positive area to the total ROI area.

### Statistical Analysis

Statistical analyses were performed on GraphPad PRISM software (Boston, MA, USA), expressed as means +/− standard error of means (SEM). Unpaired *t*-tests and ANOVA tests were used.

## Supplementary Materials

**Figure S1:** Tissue optical cleared E15 *Prx1; Ai14^f/+^* mice (control) and *Prx1-Rbm8a^f/+^; Ai14^f/+^* embryos. **A.** Lightsheet imaging of tissue optical cleared E15 *Prx1; Ai14^f/+^* (control) and *Prx1-Rbm8a^f/+^; Ai14^f/+^* embryos showing Ai14 (tdTomato) reporter signal. **B.** Representative 3D rendering images of tissue optical cleared E15 *Prx1; Ai14^f/+^* (control) and *Prx1-Rbm8a^f/+^; Ai14^f/+^* embryos. Scale bar = 100 µm. **C.** Quantification and statistical analysis of area-normalized Ai14 signal intensity in the forelimbs of E15 *Prx1; Ai14^f/+^* (control) and *Prx1-Rbm8a^f/+^; Ai14^f/+^* embryos. Ai14 signal intensity in the forelimbs was significantly reduced in *Prx1-Rbm8a^f/+^; Ai14^f/+^* embryos compared to controls. Mean ± SEM (n=4 per group). Unpaired *t*-test. **, p < 0.01. **D.** Quantification and statistical analysis of area-normalized Ai14 signal intensity in the crania of E15 *Prx1; Ai14^f/+^* (control) and *Prx1-Rbm8a^f/+^; Ai14^f/+^* embryos. Ai14 signal intensity in the crania was noticeably reduced in *Prx1-Rbm8a^f/+^; Ai14^f/+^* embryos compared to controls. Mean ± SEM (n=2 per group).

**Figure S2:** Additional quantification of craniofacial landmarks of *Prx1-Rbm8a^f/+^* and control mice across developmental timepoints. Quantification and statistical analyses of (A) frontal bone widths, (B) frontal bone lengths, (C) distance between the anterolateral corners of the frontal bone, (D) height of cranial cavity between bregma and intersphenoidal synchondrosis, (E) orbital socket width, (F) orbital socket length, and (G) representative micro-CT scan image of incisor length (left) and quantification and statistical analyses (right) of incisor length of *Prx1-Rbm8a^f/+^* mice (red) and control mice (black) across developmental timepoints (P5, P15, P30). Mean ± SEM (n=3 per group). Unpaired *t*-test. *, p < 0.05, **, p < 0.01, ***, p < 0.001, ****, p < 0.0001.

**Figure S3:** Additional results from behavioral tests on *Prx1-Rbm8a^f/+^* and control mice. **A.** Open field test results. *Prx1-Rbm8a^f/+^* mice and control mice had comparable average velocity, latency to the enter, and frequency in the center. Mean ± SEM (n=30 total; control: n=18 (9 males, 9 females); mutant: n=12 (5 males, 7 females)). Unpaired *t*-test. ns, not significant. **B.** Elevated plus maze test results. *Prx1-Rbm8a^f/+^* mice and control mice had comparable average velocity, latency to the open arms, and frequency in the open arms. Mean ± SEM (n=30 total; control: n=18 (9 males, 9 females); mutant: n=12 (5 males, 7 females)). Unpaired *t*-test. ns, not significant. **C.** Forced swimming test to measure depressive-like behavior. *Prx1-Rbm8a^f/+^* mice and control mice had comparable immobility time. Mean ± SEM (n=30 total; control: n=18 (9 males, 9 females); mutant: n=12 (5 males, 7 females)). Unpaired *t*-test. ns, not significant. Created with BioRender.com. **D.** Nesting test to evaluate general health, innate goal-directed behavior, and motor function deficits. Nest scores were recorded at 1, 3, 6, 24, and 36 hours following the introduction of nesting material. *Prx1-Rbm8a^f/+^* mice (red) had significantly lower nesting scores at earlier timepoints (3 and 6 hours) but ultimately reached comparable scores, indicating delayed nest-building behavior compared to controls (black). Mean ± SEM (n=30 total; control: n=18 (9 males, 9 females); mutant: n=12 (5 males, 7 females)). Unpaired *t*-test. *, p < 0.05, **, p < 0.01. **E.** Cylinder test to evaluate forelimb use asymmetry during vertical exploration. Both *Prx1-Rbm8a^f/+^*mice and control mice had symmetric use of left and right forelimbs. Mean ± SEM (n=30 total; control: n=18 (9 males, 9 females); mutant: n=12 (5 males, 7 females)). Unpaired *t*-test. ns, not significant.

**Figure S4:** Transcript feature analysis and pathway enrichment of RBM8A-associated RNAs reveal preferential binding to long, complex transcripts and cilia-related gene networks. **A.** Transcript feature distribution plots show that RBM8A-associated transcripts (“list”; blue) were significantly enriched for a narrower and shorter 3’ UTR length range, and a modest but significant shift toward higher GC content compared to background (“genome”; red). No significant difference was observed in 5’ UTR length between RBM8A-associated transcripts and background. Chi-squared test. **B.** Distribution of transcript biotypes for RBM8A-associated transcripts (“list”; red) compared to the genomic background (“expected”; gray). RBM8A-associated transcripts were strongly enriched for protein-coding genes. (P = 0). **C.** Distribution of number of exons per coding gene for RBM8A-associated transcripts (“list”; red compared to background (“expected”; gray). RBM8A-associated genes were significantly enriched for genes with more exons. Chi-squared test (P = 8 × 10⁻²⁵²). **D.** Chromosomal distribution plot of RBM8A-associated genes across the genome, indicating broad genomic representation without strong chromosomal bias. **E.** Heat map of transcripts enriched in RBM8A immunoprecipitates associated with RNA splicing and processing pathways showing relative expression across samples. Values represent normalized expressions and are displayed as row-wise z-scores. 3 replicates of IgG (top) on the left and 3 replicates from RBM8A immunoprecipitates (bottom). **F.** Overlap of enriched pathways identified across GO, KEGG, and Reactome databases, highlighting shared enrichment in biological processes related to the Hedgehog signaling pathway. **G.** qRT-PCR results of additional genes involved in developmental signaling pathways using RNA extracted from E15 forelimbs of *Prx1-Rbm8a^f/+^* and control embryos. Relative gene expression was quantified using the ΔΔCt method and normalized to Actin. Not statistically significant, but a trend towards reduced expression of *Grem1* and *Runx2* was observed in *Prx1-Rbm8a^f/+^* relative to controls. *Foxf1* and *Ccnd1* expressions were comparable in *Prx1-Rbm8a^f/+^* relative to controls. Mean ± SEM (n=3 per group). Unpaired t-test. ns, not significant.

**Figure S5:** Characterization of additional organs of *Prx1-Rbm8a^f/+^* mice. **A.** CBC results of adult *Prx1-Rbm8a^f/+^* (gray bar) and control mice (white bar). No statistical difference was observed in numbers of platelets (PLT), red blood cells (RBC), white blood cells (WBC), lymphocytes (LYMPH), neutrophils (NEUT), or monocytes (MONO) in blood samples of *Prx1-Rbm8a^f/+^* mice compared to those of control mice. Mean ± SEM (n=10 per group). Unpaired *t*-test. ns, not significant. **B.** Representative images of livers, hearts, kidneys, and spleens of *Rbm8a^f/+^* mice (left) and *Prx1-Rbm8a^f/+^* mice (center and right). Scale bar = 1 cm.

**Figure S6:** Generation of a platelet-specific cKO mouse model. **B.** Schematic of the breeding strategy used to generate *Pf4-Rbm8a^f/+^* mice. *Rbm8a^f/f^* mice were crossed with *Prx1-Cre* mice to generate *Pf4-Cre; Rbm8a^f/+^*offspring, which have selective deletion of *Rbm8a* in megakaryocytes and platelets. Created with BioRender.com. **C.** CBC results of adult heterozygous *Pf4-Rbm8a^f/+^* mice (solid purple bar), homozygous *Pf4-Rbm8a^f/f^* mice (patterned purple bar), and control mice (white bar). Platelet (PLT) count was significantly reduced in heterozygous *Pf4-Rbm8a^f/+^* mice, and even more drastically reduced in homozygous *Pf4-Rbm8a^f/f^* mice, compared to that of control mice. No statistical difference was observed in red blood cells (RBC), white blood cells (WBC), lymphocytes (LYMPH), neutrophils (NEUT), or monocytes (MONO) in blood samples of *Pf4-Rbm8a^f/+^* or *Pf4-Rbm8a^f/f^* mice compared to those of control mice. Mean ± SEM (n =6 per group). Unpaired *t*-test. ns, not significant, **, p < 0.01, ****, p < 0.0001. **D.** Representative images of livers, hearts, kidneys, and spleens of *Rbm8a^f/+^* mice (left), heterozygous *Pf4-Rbm8a^f/+^* mice (center), and homozygous *Pf4-Rbm8a^f/f^* mice (right). The spleen of homozygous *Pf4-Rbm8a^f/f^* mouse is noticeably larger than that of *Rbm8a^f/+^* or heterozygous *Pf4-Rbm8a^f/+^* mouse. Scale bar = 1 cm.

## Acknowledgements

We thank all members of the Mao lab for helpful discussions and support. We thank Janine Kwapis for valuable insights and constructive feedback that contributed to the interpretation of this work. We thank Thomas Neuberger and Sean Gullette at the Huck High-Field Magnetic Resonance Imaging Facility for their technical support for the micro-CT. We thank the animal facility staff for assistance with animal care.

## Author Contributions

**conceptualization**, J.M., Z.P., and Y.M.; **investigation**, J.M., C.L., Z.S., N.B., M.T., Y.K., J.Z., and Y.M.; **formal analysis**, J.M., J.S., S.G., T.P., and Y.M.; **resources**, A.V., A.L., and Y.W.; **writing - original draft**, J.M. and Y.M.; **writing - review and editing**, J.M. and Y.M.; **supervision**, Z.P. and Y.M.; **funding acquisition**, J.M. and Y.M. All authors have read and agreed to the published version of the manuscript.

## Data availability

All data necessary to support the conclusion of this study are available in the manuscript or the supplementary information. Raw data are deposited in GEO (Gene Expression Omnibus) associated with BioProject accession number PRJNA1401423.

