## Supplementary Figures for "TAR syndrome causal gene *RBM8A* is critical for embryonic bone development and proper Hedgehog signaling"

### Supplementary Figure 1

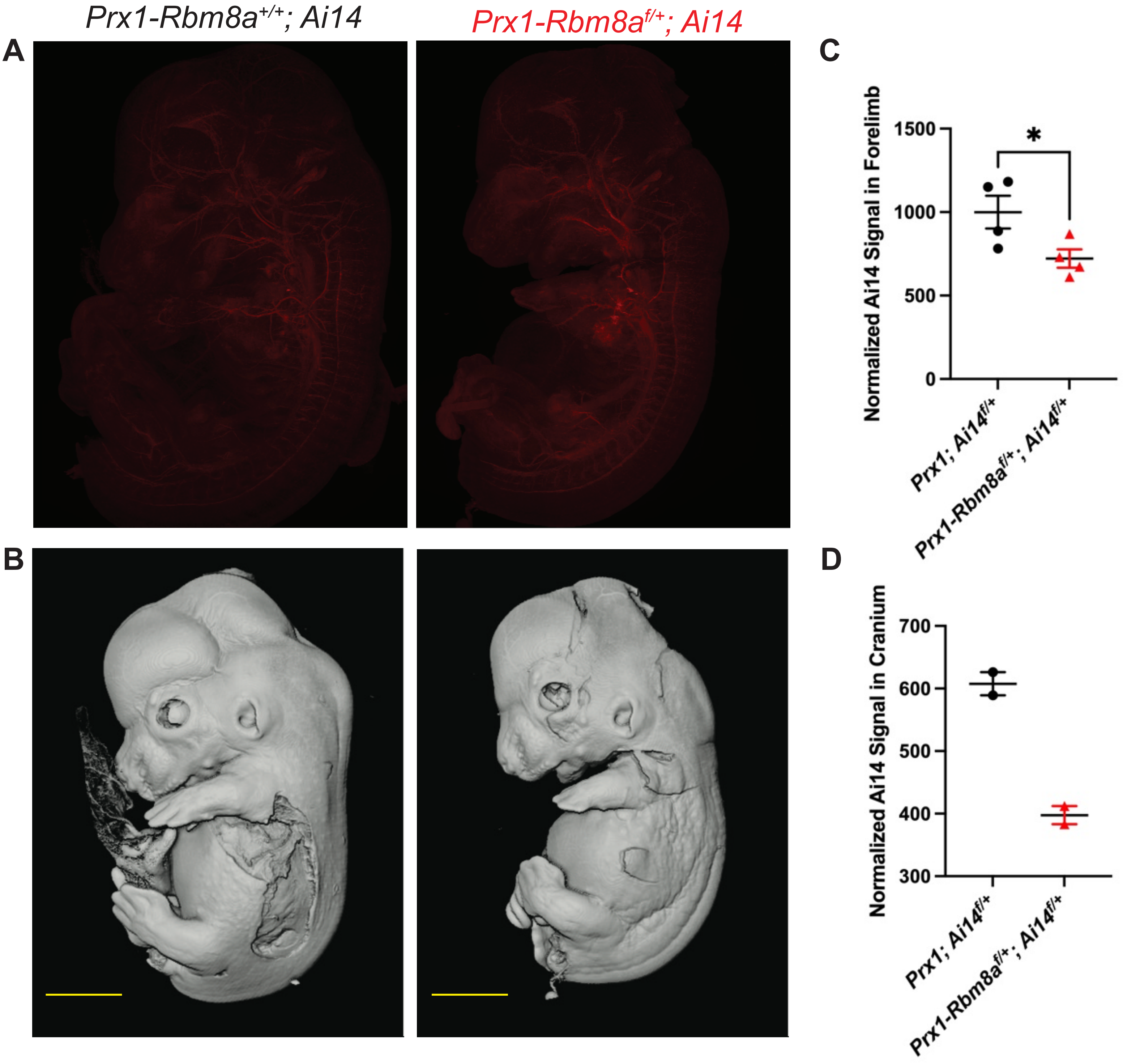

#### Supplementary Figure 2

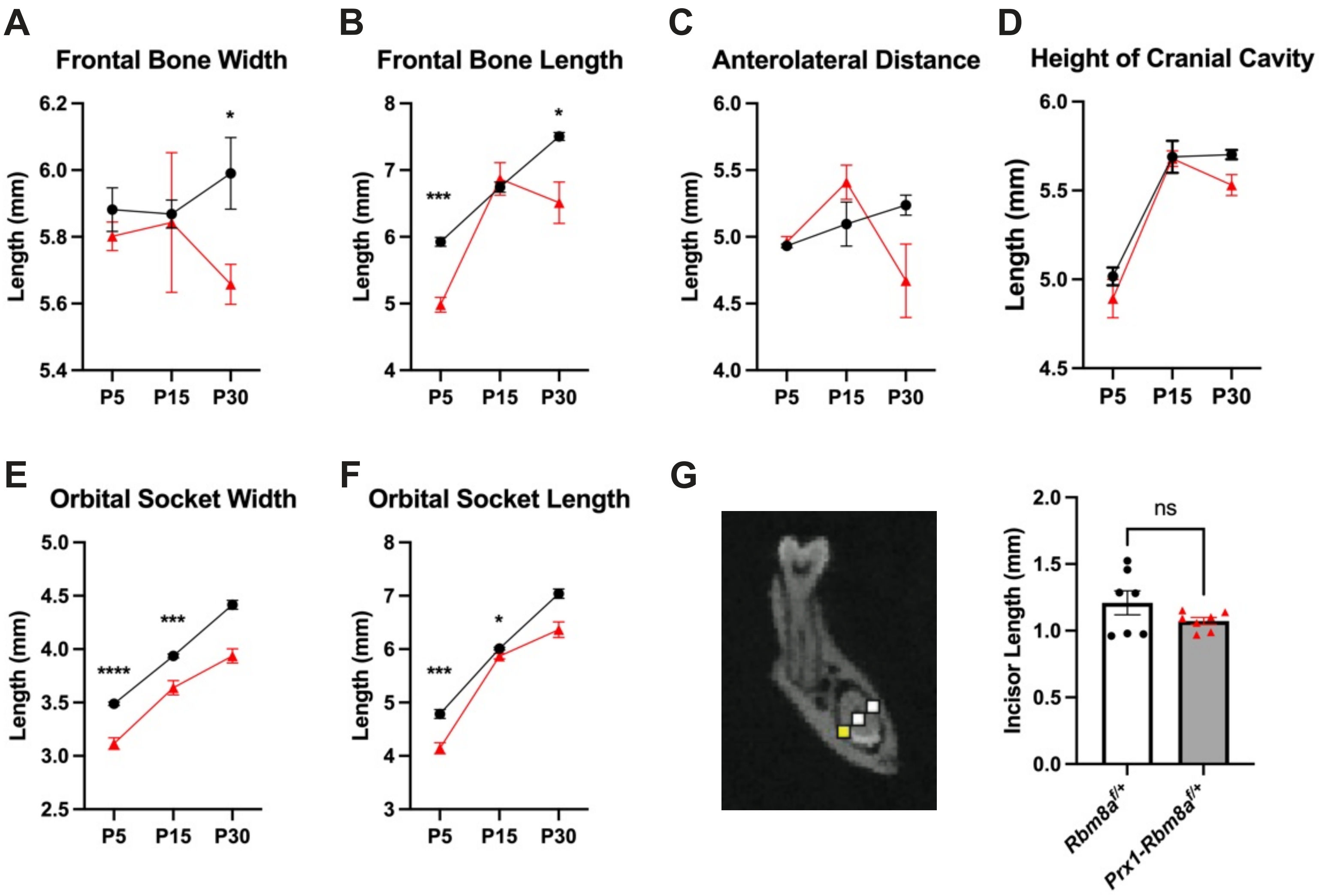

#### Supplementary Figure 3

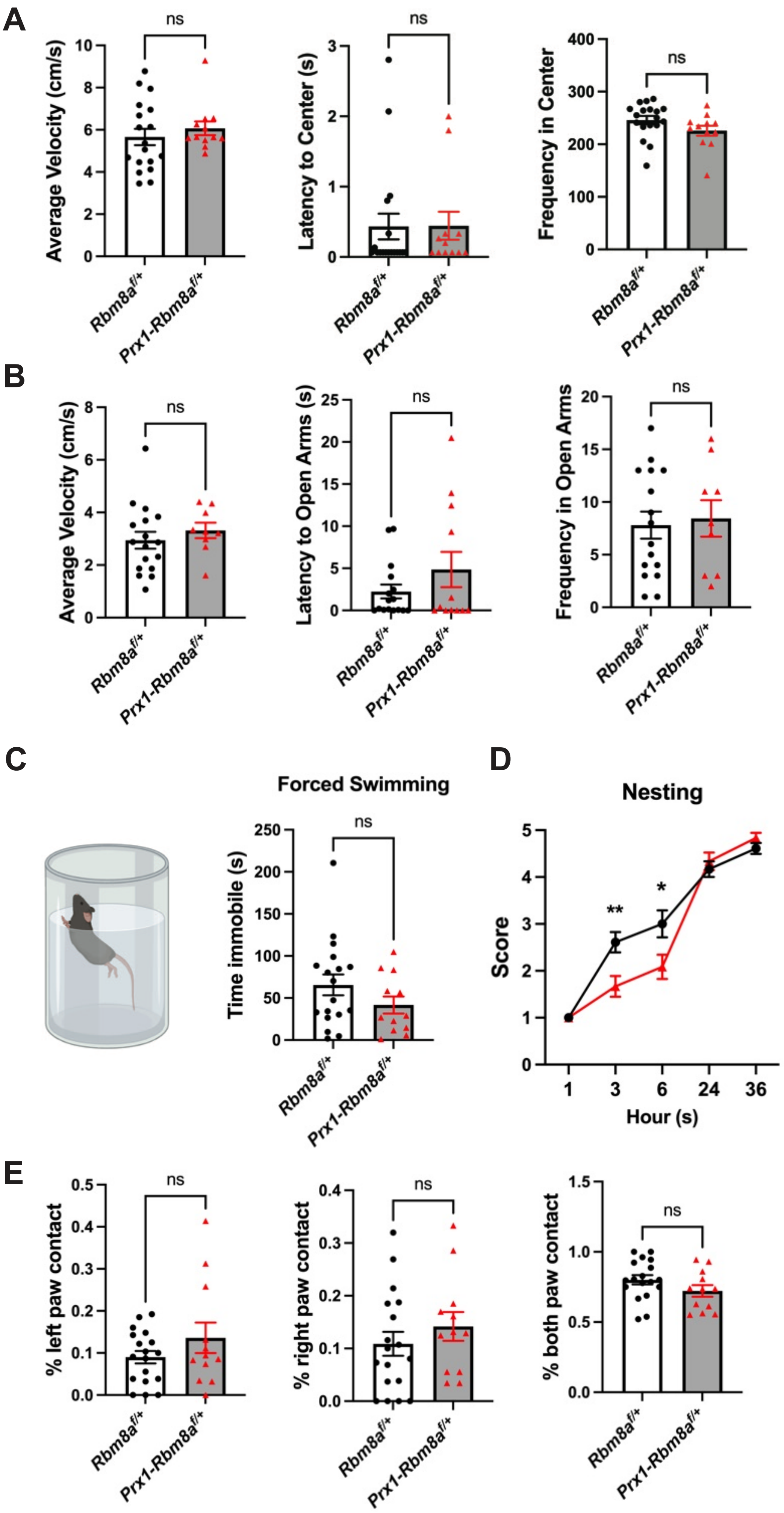

### Supplementary Figure 4

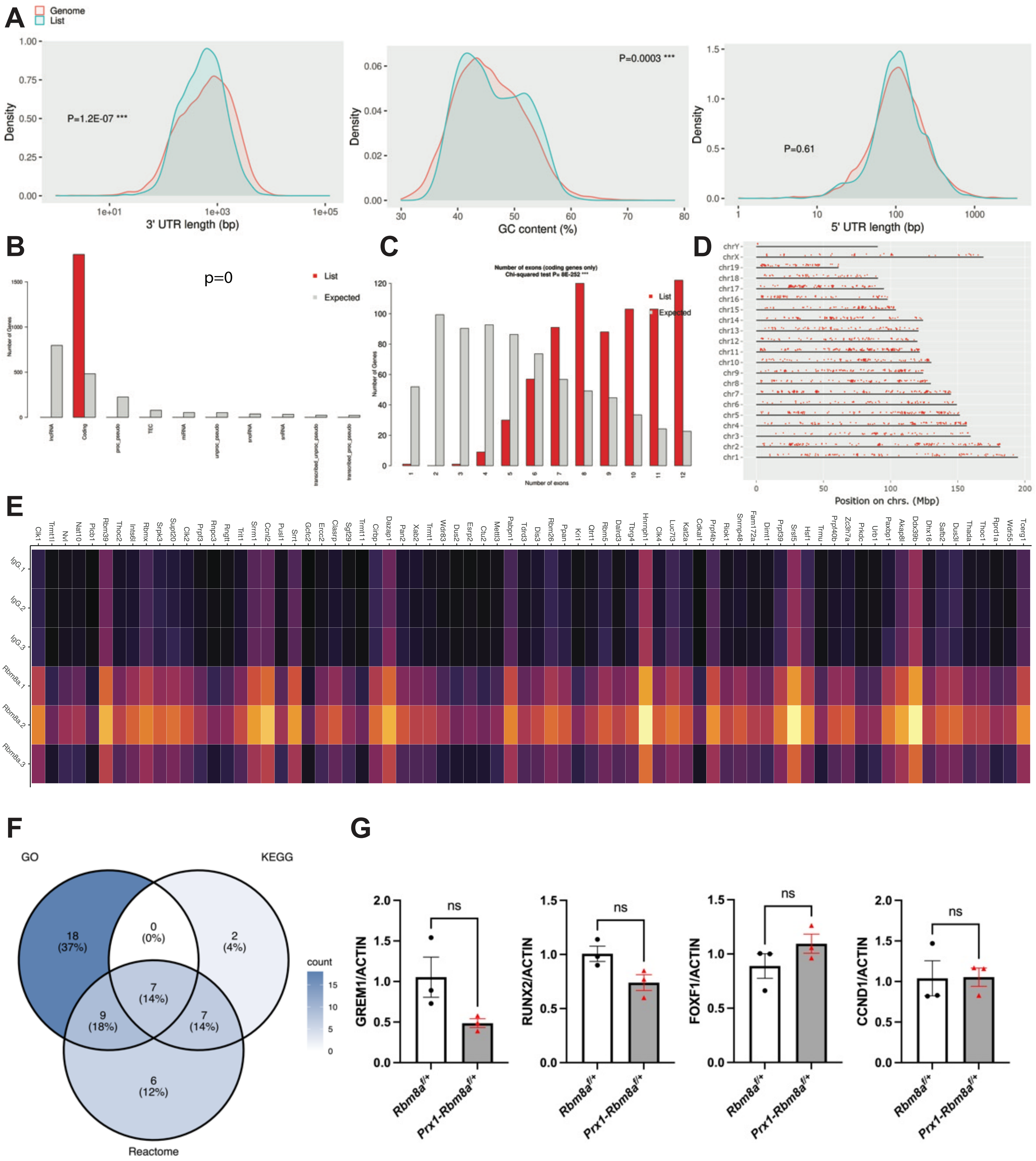

#### Supplementary Figure 5

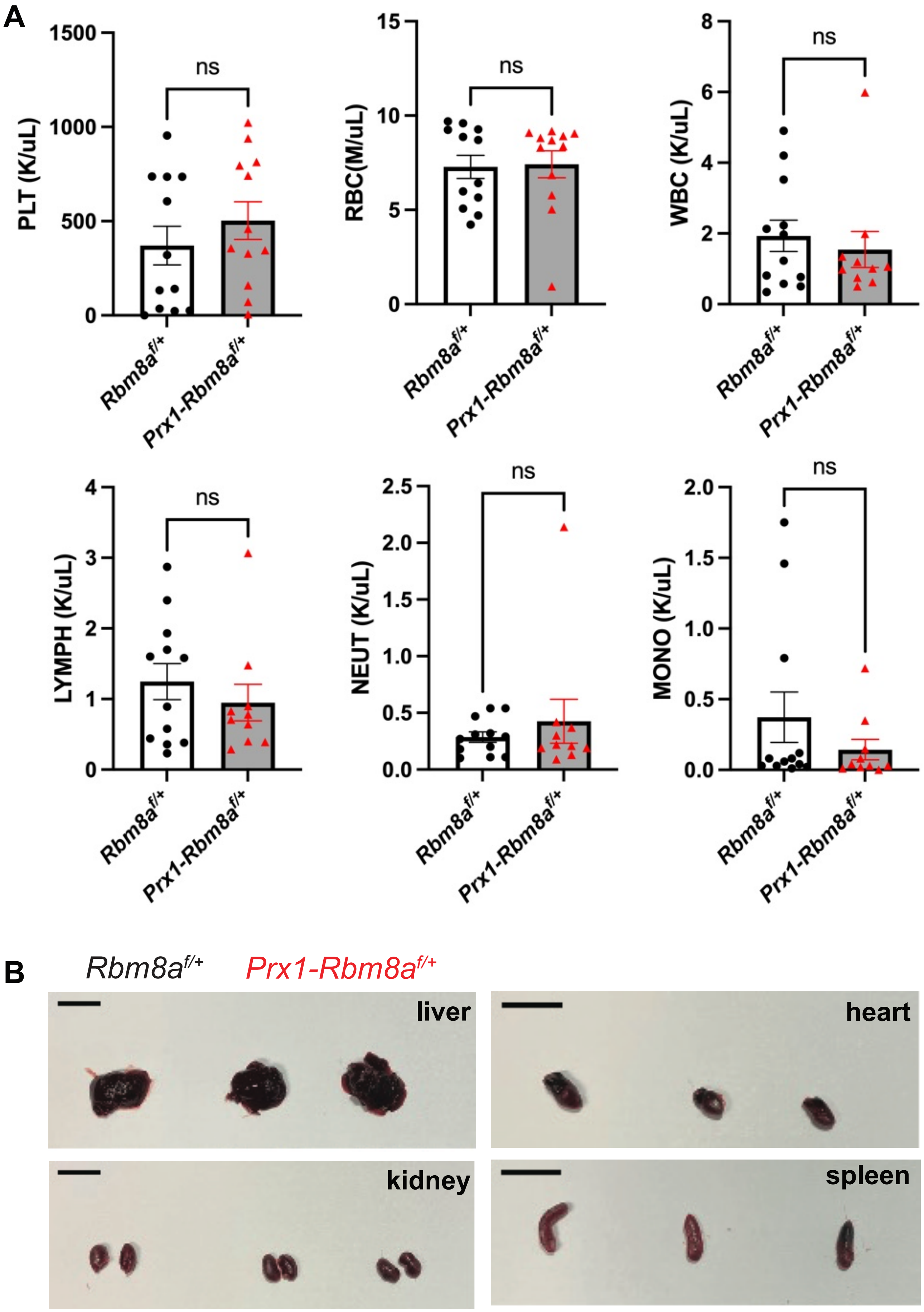

#### Supplementary Figure 6

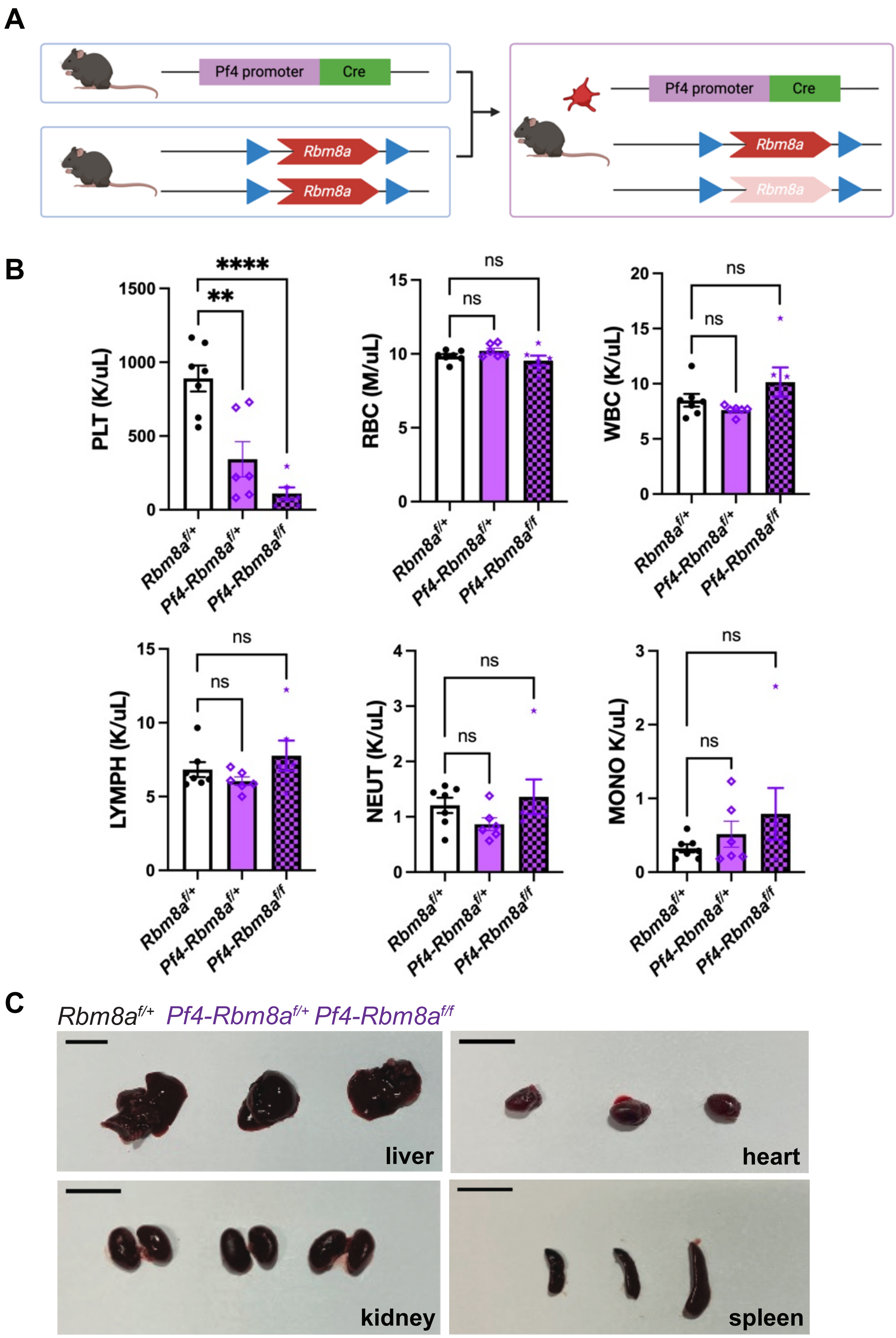
